# The VAMP associated protein, VAP27-4, negatively regulates the KAT1 inward-rectifying K^+^ channel by promoting its endocytosis, reducing guard turgor and stomatal opening

**DOI:** 10.64898/2026.11.16.767000

**Authors:** Nuria Andrés-Colás, Manuel Nieves-Cordones, Joseph Zailaa, Elisa Rodríguez-Castro, Víctor Gaya, Inés Morales-Mena, Laura Gutiérrez-Ravé, Jorge Armero-Gimenez, Laura Coloma, Lucía Torregrosa-Barragán, Pablo Coloma-Borras, Eva Montoliu-Silvestre, Borja Belda-Palazón, Jose Ramos, Craig Brodersen, José Miguel Mulet, Lynne Yenush

**Author notes:** Corresponding Author for Contact details, Nuria Andrés-Colás and Lynne Yenush, Instituto de Biología Molecular y Celular de Plantas, Universitat Politècnica de València-Consejo Superior de Investigaciones Científicas, 46022 Valencia, SPAIN.

## Abstract

Under drought conditions, plants conserve water largely by closing stomata, thereby reducing transpiration at the cost of limiting CO₂ uptake for photosynthesis. Stomatal opening and closing are driven by changes in guard cell turgor, which depend on fluxes of solutes (K⁺, anions, organic osmolytes) and water. The KAT1 inward-rectifying K^+^ channel is an important player in determining stomatal movement dynamics and so understanding its regulation and trafficking may contribute to the development of biotechnological tools for breeding or generating drought-resistant plants. To find regulators, we performed a protein-protein interaction screening in yeast. Here, we present the characterization of VAP27-4 from *Arabidopsis thaliana*, which belongs to the VAP family of proteins associated with VAMPs (vesicle-associated membrane proteins), whose function has not yet been characterized. We confirmed the KAT1-VAP27-4 interaction in planta and show that VAP27-4 acts as a negative regulator of KAT1 in both yeast and *Xenopus laevis* oocytes. We provide data showing that this putative VAP-type KAT1 regulator has a role in abscisic acid (ABA)-mediated KAT1 endocytosis, consistent with its role as a negative regulator. Importantly, we show the physiological effects of VAP27-4 co-expression on KAT1 function in guard cell turgor and stomatal opening dynamics. Our results support a role for VAP27-4 as a negative regulator of KAT1 involved in ABA-mediated channel endocytosis.

## Introduction

### Stomata and the importance of ion transport

Stomata are specialized epidermal pores that regulate the exchange of carbon dioxide and water vapor between plants and the atmosphere, playing fundamental roles in photosynthesis, transpiration, water-use efficiency (WUE), and adaptation to fluctuating environmental conditions (Assmann 1993; Blatt 2000; Schroeder et al. 2001; Lawson and Blatt 2014). Because stomatal movements simultaneously control carbon acquisition and transpirational water loss, stomatal aperture represents a major determinant of plant productivity and water balance from the cellular to the ecosystem scale (Farquhar and Cowan 1974; Lawson and Blatt 2014; Jezek and Blatt 2017). Changes in stomatal aperture are driven by reversible alterations in guard-cell turgor that arise from coordinated fluxes of K⁺, anions, organic osmolytes, and water across the plasma membrane (PM) and endomembrane system (Blatt 2000; Meckel et al. 2007; Lawson and Blatt 2014; Jezek and Blatt 2017). H⁺-ATPases generate pH and voltage gradients that energize the transport of osmotically active solutes, principally K⁺, Cl⁻, and malate, thereby driving water uptake and changes in guard-cell volume (Schroeder et al. 1987; Blatt 1992, 2000, 2024). Among these transport processes, K⁺ uptake through inward-rectifying Voltage-gated K⁺ channels, particularly KAT1 and KAT2, constitutes a major determinant of stomatal opening (Nakamura et al. 1995; Ichida et al. 1997; Blatt 2000; Pilot et al. 2001; Lebaudy et al. 2008; Jezek and Blatt 2017). Conversely, stomatal closure requires coordinated activation of outward K⁺ and anion efflux pathways, including the guard-cell outward rectifier GORK and the S-type anion channels SLAC1 and SLAH3, which drive the loss of osmotically active solutes and promote guard-cell shrinkage (Ache et al. 2000; Hosy et al. 2003; Negi et al. 2008; Zhang et al. 2016, 2025). The activities of these ion channels and plasma membrane H⁺-ATPases are highly regulated, underpinning stomatal responses to light, the stress hormone abscisic acid (ABA), atmospheric CO₂, and other environmental signals (Blatt 2000; Negi et al. 2008; Assmann and Jegla 2016).

### KAT1

The inward-rectifying K⁺ channel, KAT1, was one of the first ion transport proteins cloned and functionally characterized (Anderson et al. 1992; Schachtman et al.

1992). It is expressed almost exclusively in guard cells and encodes a channel that is activated by plasma membrane hyperpolarization, mediating K⁺ uptake during stomatal opening (Nakamura et al. 1995; Ichida et al. 1997). Genetic analyses suggest that additional channels are involved, as inward K⁺ currents and guard-cell function remain in *kat1* mutants (Szyroki et al. 2001). This observation established that KAT1 operates within a broader network of inward-rectifying K⁺ channels, including KAT2, AKT1, AKT2 and KC1 (Pilot et al. 2001; Szyroki et al. 2001). Two different functional studies in transgenic *A. thaliana* lines showed that dominant-negative mutations in either *KAT1* or *KAT2* significantly reduce inward K⁺ currents, which results in a decrease in light-induced stomatal opening, and a concomitant reduction in water loss through transpiration (Kwak et al. 2001; Lebaudy et al. 2008). Further evidence supporting the importance of K^+^ fluxes to improve WUE without a penalty in carbon fixation was provided in a study where a synthetic, light-gated K+ channel, BLINK1, was expressed in guard cells and shown to accelerate both stomatal opening and closure and drive a more than 2-fold increase in biomass in fluctuating light (Papanatsiou et al. 2019). These studies provide support for the vital role of K^+^ channels, and particularly KAT1, in stomatal movements and therefore, the response of plants to water deficit.

KAT1 activity is controlled by multiple signaling pathways. Early electrophysiological studies demonstrated that inward K⁺ channels are strongly regulated by cytosolic Ca²⁺ and pH, two central intracellular signals associated with environmental and hormonal responses (Blatt and Grabov 1997; Blatt 2000). This channel is integrated into guard-cell signaling through direct interactions with anion channels. The S-type anion channel SLAC1 and the related channel SLAH3 play central roles in ABA-induced stomatal closure and CO₂ responses (Negi et al. 2008; Zhang et al. 2016). Importantly, SLAC1 physically associates with KAT1 and strongly inhibits KAT1-mediated inward currents, whereas SLAH3 exerts a more modest effect (Zhang et al. 2016). A particularly important regulatory pathway affecting KAT1 involves ABA, the central phytohormone controlling stomatal closure during water deficit. ABA signaling is closely associated with elevations in cytosolic Ca²⁺ that suppress inward K⁺ transport and plasma membrane H⁺-ATPase activity while promoting activation of anion efflux pathways, thereby coordinating the transition from stomatal opening to closure (Assmann and Jegla 2016; Jezek and Blatt 2017). Activation of the ABA-responsive SnRK2 kinase OPEN STOMATA1 (OST1/SnRK2.6) is required for key components of guard-cell ABA signaling, including suppression of inward K⁺ currents and activation of anion fluxes (Acharya et al. 2013). OST1 physically interacts with KAT1 and phosphorylates residues within its cytoplasmic domain (Sato et al. 2009; Acharya et al. 2013).

### KAT1 trafficking-importance and known regulators

Increasing evidence indicates that stomatal signaling is regulated not only through modulation of transporter activity but also through dynamic changes in transporter abundance at the plasma membrane (Eisenach et al., 2012a; Jezek & Blatt, 2017; Karnik et al., 2017a). Vesicle trafficking selectively inserts and retrieves membrane proteins, thereby altering guard-cell transport capacity and influencing the kinetics and magnitude of stomatal responses (Xue et al. 2018; Xia et al. 2019; Baena et al. 2022, 2024).

Newly synthesized KAT1 channels enter the endoplasmic reticulum (ER) and travel through the secretory pathway before reaching the cell surface. A conserved diacidic motif within KAT1 is required for efficient export from the ER and interacts with the COPII cargo-selection factor Sec24 (Mikosch et al. 2006; Sieben et al. 2008; Reuff et al. 2010). BAG proteins function in cellular protein quality-control pathways, and BAG4 has been shown to physically interact with KAT1, enhance channel activity, and promote its accumulation at the plasma membrane (Locascio et al. 2019).

Following delivery to the PM, KAT1 remains dynamically organized rather than freely diffusing throughout the membrane accumulating within relatively stable PM microdomains (Sutter et al. 2006). A major regulator of this organization is the Qa-SNARE SYP121, a member of the conserved SNARE family of membrane-fusion proteins (Söllner and Rothman 1996; Jahn and Südhof 1999; Jürgens 2004; Bassham and Blatt 2008). SYP121 participates in vesicle fusion at the PM and influences KAT1 localization (Sutter et al. 2006). Beyond its trafficking role, SYP121 directly interacts with KAT1 and modulates channel gating through a persistent physical interaction that alters channel energetics and promotes channel opening when the SNARE is anchored at the PM (Lefoulon et al. 2018). This interaction involves a conserved FxRF motif found in several Voltage-Gated K^+^ channel family members and couples channel gating to membrane trafficking processes (Honsbein et al. 2009, 2011). Subsequent studies demonstrated that SYP121 can directly engage K⁺-channel voltage-sensing machinery, thereby coordinating membrane transport, secretory traffic, cell turgor regulation, and growth (Grefen et al. 2015; Karnik et al. 2017; Lefoulon et al. 2018).

The R-SNARE VAMP721 protein also interacts directly with KAT1, reducing channel activity at the PM (Zhang et al. 2015). Simultaneously, the microtubule plus-end-binding protein EB1b directly interacts with SYP121 and supports a previously unrecognized trafficking pathway linked to microtubule growth toward the PM (Zhong et al. 2024). Disruption of this pathway alters the distribution of both SYP121 and KAT1 and delays stomatal opening (Zhong et al. 2024).

The converse process, endocytosis, is equally important for regulating channel abundance. and guard-cell transport capacity and stomatal behavior (Eisenach et al. 2012, 2014; Karnik et al. 2017). ABA promotes selective endocytosis of KAT1 from the PM, leading to accumulation of the channel within intracellular endosomal compartments (Sutter et al. 2007). Following removal of ABA, the channel gradually recycles back to the PM over several hours via a SYP121-dependent route (Sutter et al. 2007; Eisenach et al. 2012) Studies of both KAT1 and GORK further indicate that ion channels can form dynamic membrane clusters whose organization is intimately connected to channel regulation and membrane trafficking processes (Sutter et al. 2006; Eisenach et al. 2014). Recent work has extended this model to environmental regulation. RPT2 physically interacts with KAT1, promotes KAT1-mediated currents, and contributes to channel accumulation (Locascio et al. 2025). Moreover, elevated atmospheric CO₂ alters KAT1 mobility at the PM and stimulates channel internalization via a process that depends on SYP121 (Yu et al. 2026).

### ER-PM contact sites and VAP27 family proteins

A growing body of work indicates that membrane trafficking is itself spatially organized by membrane contact sites (MCSs), such as ER-PM contact sites (EPCSs), which coordinate lipid exchange, signaling, cytoskeletal organization, and membrane trafficking (Wang et al. 2017). Among the principal organizers of these structures are the VAP27 proteins, a family of ER-resident tethering proteins homologous to the yeast Scs2/VAP proteins (Wang et al. 2016). These proteins contain an MSP-domain, an N-terminal CCD-domain, and a C-terminal TMD domain, for anchoring to the ER (Wang et al. 2016). VAP27 proteins localize to EPCSs throughout the plant and participate in the formation of stable ER-PM connections that are closely associated with both the actin and microtubules (Wang et al. 2016, 2017).

Arabidopsis contains ten VAP27 family members, with VAP27-1, VAP27-3, and VAP27-4 among the best-characterized isoforms at EPCSs (Wang et al. 2016). Importantly, genetic analyses demonstrated that VAP27 proteins are required for normal plant growth and development, with simultaneous disruption of multiple VAP27 isoforms leading to severe defects in root growth and ER-PM connectivity (Wang et al. 2016). Particularly relevant to membrane trafficking, VAP27-1 and VAP27-3 interact with clathrin and phosphoinositides enriched endocytic membranes and are required for normal clathrin-mediated endocytosis (CME) (Stefano et al. 2018). VAP27-dependent contact sites therefore represent candidate spatial regulators that couple endocytosis to broader aspects of membrane organization and signaling. More recent studies have further linked VAP27 proteins to aquaporin trafficking, autophagy-related proteins, exocyst components, organelle communication, and stress adaptation, supporting an emerging view in which VAP27 proteins function as multifunctional organizers of membrane dynamics and inter-organelle communication rather than static ER-PM tethers (Fox et al. 2020; Codjoe et al. 2022; Li et al. 2022; Zhang et al. 2023; Man et al. 2024).

Despite considerable progress in elucidating guard-cell signaling networks, the mechanisms that regulate the trafficking and turnover of key ion channels remain poorly understood. A major outstanding question concerns the molecular machinery responsible for KAT1 endocytosis and its integration into stomatal signaling pathways. In this study, we identify VAP27-4 as a previously unrecognized KAT1-interacting protein that promotes KAT1 endocytosis and modulates stomatal opening dynamics. These findings shed light on the KAT1-endocytosis machinery and thereby provide a framework for understanding how guard cells translate environmental and hormonal cues into dynamic changes in membrane protein composition, ion fluxes, and stomatal aperture.

## Materials and Methods

### Yeast split-ubiquitin and functional complementation assays

Split-Ub protein-protein interactions assays were performed as described (Locascio et al. 2019) using the KAT1-pMetYCgate vector (ABRC CD3-815; (Obrdlik et al. 2004)) co-transformed with the pDSL-Nx vector containing the *VAP27-4* clone recovered in the screening (lacking the first 15 amino acids of the VAP27-4 (AT5G47180) protein) into the THY.AP4 *Saccharomyces cerevisiae* strain (MATa, *ura3*, *leu2*, lexA::lacZ::*trp1*, lexA::*HIS3*, lexA::*ADE2*) (Paumi et al. 2007). For the functional complementation assays, these vectors were co-transformed into the PLY240 strain (MATa *his3*Δ200 *leu2*-3,112 *trp1*Δ901 *ura3*-52 *suc2*Δ9 *trk1*Δ51 *trk2*Δ50::lox-*KanMX*-lox; (Bertl et al. 2003) and for growth in solid media, transformants were grown to saturation in SD media supplemented with 100 mM KCl and the cells were centrifuged and washed with sterile ultrapure water at 8000 rpm and 4 °C for 5 minutes, after which the cell numbers were adjusted to A600nm = 1. Serial dilutions were plated onto Translucent media (YNB-F; ForMedium™ UK; (Navarrete et al. 2010)) with the necessary auxotrophic requirements and supplemented with 2 % agar and limiting concentrations of KCl or toxic concentrations of LiCl, as indicated. Growth was recorded after 72 hours of growth at 28°C. For growth assays in liquid media, cells from saturated cultures washed with sterile ultrapure water (8000 rpm, 4 °C, 5 minutes) and adjusted to A600nm = 1. These cell suspensions were then inoculated into tubes containing 5 mL of YNB-F medium supplemented with the necessary auxotrophic requirements and different concentrations of KCl (2 mM and 20 mM) at A600nm = 0.05. The inoculated tubes were incubated at 28 °C under agitation (170 rpm). Absorbance was measured at the indicated times using a Spectronic 20D (Milton Roy) spectrophotometer.

### Plant Materials and Growth Conditions

The following *Arabidopsis thaliana* lines were employed: WT (Col-0), *kat1-2* (SALK 093506, T-DNA insertion in the first exon (Nozue et al., 2015)) and *vap27-4* (SAIL_1255 D12 / N866874 from the Nottingham Arabidopsis Stock Centre (NASC) Collection). We confirmed both the genotype (PCR) and lack of *VAP27-4* expression (RT-qPCR) (Fig. S1). The WT and KO plants were transformed using the floral dipping method (Clough and Bent 1998) with the *pDGB3 omega1* GoldenBraid plasmids containing: *KAT1-YFP, alone or with HA-VAP27-4, VAP27-4-YFP* or *YFP-VAP27-4,* both under the control of the cauliflower mosaic virus *CaMV35S* promoter and with the *T_nos_* terminator, together with *DsRed,* under control of seed expression *At2S-3* promoter, or *hph,* under control of P_nos_ promoter, for plant selection. Arabidopsis plants were grown under a 16 h light/8 h dark photoperiod at 22°C in 0.5x Murashige and Skoog media supplemented with 1 % sucrose. *Nicotiana benthamiana* plants were grown for 4–5 weeks in soil under a 16 h light/8 h dark photoperiod at 24°C. For endocytosis assays, seedlings were grown for 10 days (long day light cycle) and were treated with 100 µM ABA and 50 µM Brefeldin A for 1 h in liquid 0.5x Murashige and Skoog media supplemented with 1 % sucrose, before confocal fluorescence microscopy analysis carried out as described below.

#### RT-PCR and RT-qPCR

100 mg of the aerial part of 18-day-old plants grown in 0.5x MS media were collected and frozen in liquid N_2_. Samples were ground by introducing 2 metal beads into each tube and pulsing for 30 s at 30 Hz in a Retsch mixer mill, freezing the samples in liquid N_2_ between pulses. The RNA extraction was performed using the NucleoSpin RNA Plant kit (Macherey-Nagel). cDNA was synthesized from 500 ng of RNA using the PrimeScript RT Reagent Kit (Takara Bio), using both oligo dT and random hexamer primers. The PCR reactions were prepared with *Supreme NZYTaq II* at denaturation (15 min at 95°C), 38 cycles of amplification (15 s at 95°C, 30 s at 63.7°C and 30 s at 72°C) and elongation (10 min at 72°C). The qPCR reactions were prepared with 5X PyroTaq EvaGreen qPCR Mix Plus (ROX; Cultek). The reactions were carried out in a QuantSudio 3 (ThermoFisher) following the following program: activation of the hot start DNApol and denaturation (15 min at 95°C), 40 cycles of amplification (15 s at 95°C, 30 s at 57°C and 30 s at 72°C) and melting curve (1 min at 60°C, heating to 95°C at 0.1°C/s and 15 s at 95°C). In each sample, the GADPH gene was amplified as an endogenous control. The results were analyzed comparing the 2^-ΔΔCq^ in each case. The primers used are listed in Table S1.

#### Co-immunopreciptiation and BiFC assays

Experiments were carried out as described previously (Locascio et al. 2019). 3-4-week-old *N. benthamiana* plants were infiltrated with agrobacterium strains (C58Ci), one transformed with the plasmid encoding the P19 Tomato Bushy Stunt Virus silencing suppressor (Sarrion-Perdigones et al. 2013) and one with the corresponding GoldenBraid plasmids (Sarrion-Perdigones et al. 2013) carrying the indicated constructs under the *CaMV35S* promoter. Leave samples were taken and analyzed 72 hours after the agroinfiltration. For co-IP assays, extraction of total proteins was carried out in a buffer containing 50 mM Tris pH 7.5, 5 mM EDTA, 150 mM NaCl, 1 mM Phenylmethylsulfonyl fluoride (PMSF), 20% glycerol and 0.15% NP-40 and protease inhibitor cocktail (Roche), for 15 minutes at 4°C in a rotating mixer at its lowest speed. The material was then filtered using Whatman® folded filters previously hydrated with the same extraction buffer. The KAT1-YFP protein was purified using the GFP-trap (GFP-Trap®_MA, Chromotech) according to the manufacturer’s indications. Protein samples were analyzed as described in the Western blotting section. For BiFC assays, samples were analyzed as described in the confocal fluorescence microscopy section.

#### Western Blotting

Extraction of total proteins, except for co-IP assays, was carried out in 2.5x LSB buffer (4.75% SDS, 62.5 mM dithioeritriol, 6.25 mM EDTA, 18.75% sacarose, 0.156 mg/ml bromophenol blue, 0.19 M Tris-HCL pH 6.8). Proteins in samples were separated on 8% SDS-PAGE gels, transferred to nitrocellulose membranes. Membranes were stained with 0.008% Direct Blue (Aldrich Chemistry) and probed with anti-GFP (PABG1, ChromoTek), anti-HA (clone 12CA5, Roche, and clone 16B12, Covance Inc., mixture), anti-YFN and anti-YFC antibodies (AS11 1776 and AS11 1775, Agrisera). Immune complexes were visualized using the corresponding anti-mouse or anti-rabbit, respectively, HRP secondary antibodies and SuperBright ECL detection system (Agrisera). Chemiluminescence was recorded using an Amersham ImageQuant 800 imaging system.

#### Confocal fluorescence microscopy

The fluorescent proteins used in this study were: the yellow fluorescent protein (YFP); plasma membrane marker, AtPIP2A-mCherry in the pBIN20 vector (kindly provided by Dr. María Jesús Marcote, UV, Spain); endoplasmic reticulum marker, HDEL-ChFP, pCBNoXbaSTtmdCherry; vacuole marker, ChFP-VAMP713 and Golgi marker, pCBNoXbaSTtmdCherry (kindly provided by Dr. Vicente Pallás, IBMCP (UPV-CSIC), Spain); EPCS marker (kindly provided by Dr. M. Botella, IHSM (CSIC-UMA), Spain);and At2S-3:DsRed (kindly provided by Dr. M.D. Garcerá, IBMCP (CSIC-UPV), Spain). For the visualization of the fluorescent proteins 72 hours after the infiltration in *Nicotiana benthamiana* leaves, or in steady transformed *Arabidopsis thaliana* leaves, an inverted laser scanning confocal microscope (Zeiss LSM780) was used, using the argon laser with a Plan-Apochromat 40x/1.20 W DIC M27 objective and the ZEN 2001 software (Zeiss). YFP fluorescence was visualized by exciting at λ = 514 and emission was analyzed at a range of λ = 518-544/551 (endocytosis and localization in Arabidopsis) / 571 (localization and BiFC in Nicotiana) nm. The fluorescence of mCherry and ChFP were visualized by exciting at λ = 561 and emission at 589-642 nm. The emission peak of all fluorescent signals was confirmed. For the endocytosis assay, z-stack images of single stomata in true leaves of 10-day-old Arabidopsis seedlings grown on 0.5x MS with 1% sucrose were taken after treatment with 100 µM ABA and 50 µM BFA for 1 h in liquid 0.5x MS with 1% sucrose. The z-stacks and the single sections were analyzed using the Image J FIJI software (Schneider et al. 2012).

#### Two-electrode voltage clamp in Xenopus oocytes

KAT1, KAT2 and VAP27-4 cDNAs were amplified by PCR with modified oligonucleotides to add the T7 promoter at the 5’ end (Table S1). Then, cRNA synthesis was carried out on these samples with the mMACHINE T7 Ultra kit (Ambion; http://www.ambion.com). 1 ng of KAT1 or KAT2 and VAP27-4 cRNAs were injected as indicated in stage V-VI Xenopus oocytes using a Nanoliter 2000 microinjector (World Precision Instruments). Oocytes were kept in ND96 solution at 18 °C for 3-5 days before used in two-electrode voltage-clamp experiments. Oocyte membrane potential was clamped with an Axoclamp 900A amplifier (Molecular Devices). pClamp software (Molecular Devices) was used to record and analyze data. Recording solution contained (in mM): 100 potassium gluconate, 1 CaCl_2_, 2 MgCl_2_, and 10 2-(N-Morpholino)ethanesulfonic acid (MES),-NaOH, pH 5.5. The steady-state current intensity was used in I/V plots. Membrane voltages ranged from 0 to -130 mV (increments of 10 mV; holding potential, -20 mV; deactivation potential, -20 mV).

#### Cavitation bubble manometry (guard cell turgor estimation)

Seeds were surface sterilized prior to sowing using a 50% bleach solution including 0.05% Triton X-100 for 5 minutes in microcentrifuge tubes on a shaker tray prior to being rinsed twice with autoclaved DI water. Following surface sterilization, seeds were stratified at 4 °C for 3 days before sowing seeds in half-strength MS media composed of 2.2 g L^-1^ of MS basal salts, 50 mg L^-1^ of MES, 10 g L^-1^ of sucrose, and 8 g L^-1^ of phytoagar. The plates were placed in a growth chamber under short-day conditions (8 h light and 16 h dark; 50 - 100 μmol m^−2^ sec^−1^ of photosynthetic photon flux density; 22 °C; 70% relative humidity) for 10 days prior to measurements.

We used cavitation bubble manometry (CBM; (Brodersen et al. 2025), Zailaa et al., under review) to determine how guard cell turgor responded to light. Plants were removed from the growth chamber at three different time points for analysis: 1 hour before the start of the photoperiod (-1h timepoint), 3 hours after the start of the photoperiod (3h timepoint), and 9 hours after the start of the photoperiod (9h timepoint, i.e., 1 hour after the end of the photoperiod). For plants sampled during periods of darkness, the plates were wrapped in aluminum foil during transit from the growth chamber to the lab. The CBM system, previously described in detail ((Brodersen et al. 2025) ; Zailaa et al., under review) consists of an upright fluorescence microscope with a MicroPoint laser ablation system (Model 337-USAS, MicroPoint, Andor, Concord, Massachusetts, USA) and configured with a digital c-mount camera (model #acA640-750um, Basler, Inc., Exton, Pennsylvania, USA). Briefly, the system is used to vaporize a small volume of liquid inside the target cell resulting in the formation of a gas microbubble with a radius of <5 µm. By monitoring the growth and dissolution of the microbubble with a camera operating at ∼1,000 frames per second it is possible to observe relative changes in microbubble radius, which is inversely proportional to the pressure of the surrounding fluid. Plants were removed from the agar plate and placed inside the well of a depression slide surrounded by moist paper towels to maintain high relative humidity. We generated one microbubble in five to eleven guard cells from one to seven individuals and recorded their formation and dissolution dynamics. From each image sequence, we measured the maximum microbubble radius coinciding with the first frame in which the microbubble appears in the guard cell using imageJ (FIJI Version 2.16.0). Most (∼98%) of measurements were completed within 15 minutes of leaf tissue exposure to the transmitted light of the optical system.

#### Stomatal aperture assay

Cotyledons from 4-day-old Arabidopsis seedlings, grown under an 8 h dark/16 h light photoperiod at 23°C in 0.5x MS plates with 0% sucrose, were analyzed under a Leica 5000B (Leica Microsystems ©) microscope equipped with 20x objective and a Leica DFC 550 digital camera using Normarski or DIC technique and LAS V4.9. software. Stomata aperture ratio (width/length) was then measured for at least 100 stomata/condition using ImageJ FIJI software.

#### Statistics

The *GraphPad t Test Calculator* software was used for Unpaired t test comparing each sample to the control line for the endocytosis assays and to analyze gene expression differences. The microbubble radii data was analyzed using a two-way analysis of variance (ANOVA) using the timepoints (-1h, 3h, and 9h relative to the start of the photoperiod) and genotype as categorical effects, including their interaction. ANOVA model assumptions were assessed using residual diagnostics plots showing no departure from homoscedasticity or normality, the Shapiro-Wilk test of residual normality (p = 0.686), and Leven’s test of homogeneity of variance (p = 0.273). Because the interaction between the timepoint and genotype factor was significant, we assessed pairwise comparisons using tukey-adjusted estimated marginal means to compare timepoint data between genotypes and genotype data between timepoints. All statistical analyses were conducted using R Statistical Softwar (v4.3.1; RCore Team 2023).

## Results

### Identification of VAP27-4 as a KAT1 interactor

Given the important role of K^+^ channels, and particularly KAT1, in stomatal movements and the response of plants to water deficit, we carried out a high-throughput Split-Ub protein-protein interaction screen in yeast to look for potential KAT1 regulators. Using this screen, we identified and subsequently characterized three positive regulators of KAT1, BAG4, MEE31 and RPT2 (Locascio et al. 2019, 2025; González-García et al. 2024). In the present study, we present results for a novel KAT1 interactor, VAP27-4, which was also identified in this screen. Fig. 1 shows the confirmation of the interaction in yeast, and in Fig. 2, we present the results of the co-immunoprecipitation (Co-IP) and bimolecular fluorescence complementation (BiFC) analyses carried out in transient expression experiments in *N. benthamiana* to confirm that this interaction also takes place in plants (for additional control experiments related to the BiFC analysis, see Fig. S2). Co-localization experiments indicate that the interaction between KAT1 and VAP27-4 takes place in the ER and vacuole/vesicles but not in Golgi (Fig. S3). We observed that VAP27-4 also interacts with KAT2, (CoIP and BiFC) (Fig. S4), but not with AKT1 (BiFC) (Fig. 2).

**Figure 1.**
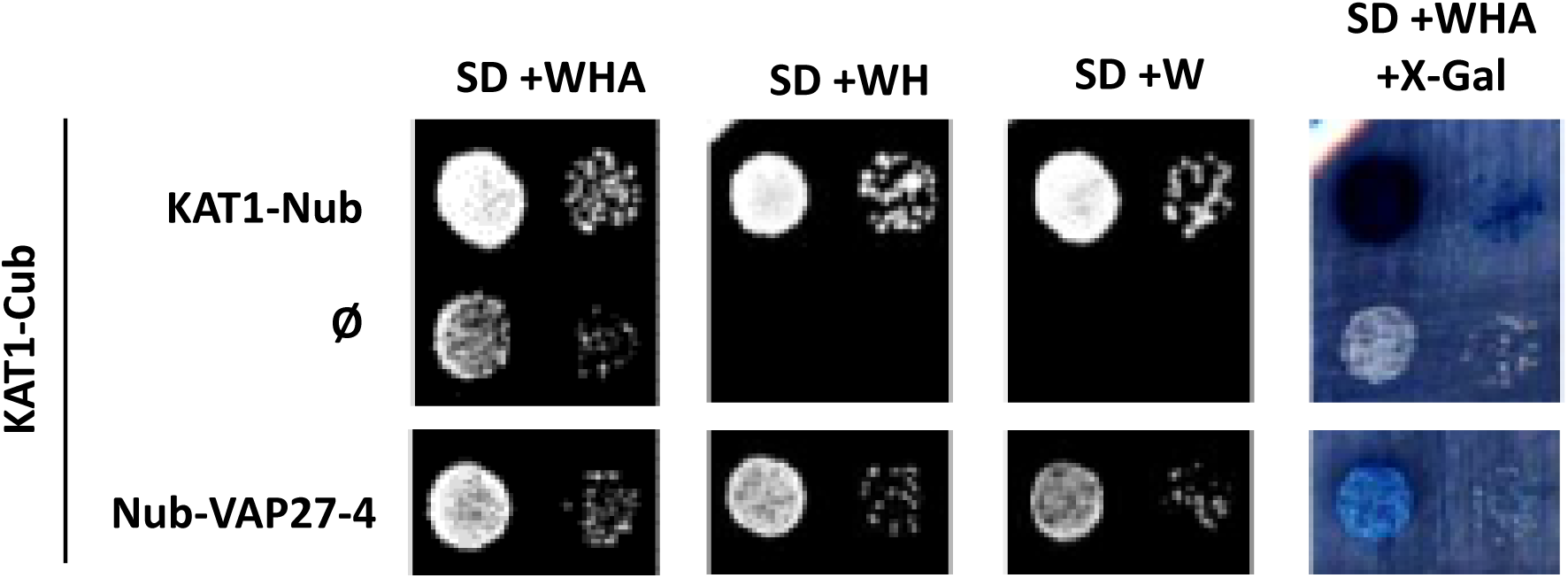
VAP27-4 and KAT1 interaction in yeast by Split-ubiquitin assay. The THY.AP4 yeast strain was co-transformed with the KAT1-Cub vector and the plasmids indicated on the left of the figure. Saturated cultures were spotted onto the indicated media, and growth was recorded after 72 hours. Similar results were observed in 4 independent clones for each combination. KAT1 was used as a positive control. Ø, empty vector. SD, Synthetic Defined Medium. W, tryptophan. A, alanine. H, histidine.

**Figure 2.**
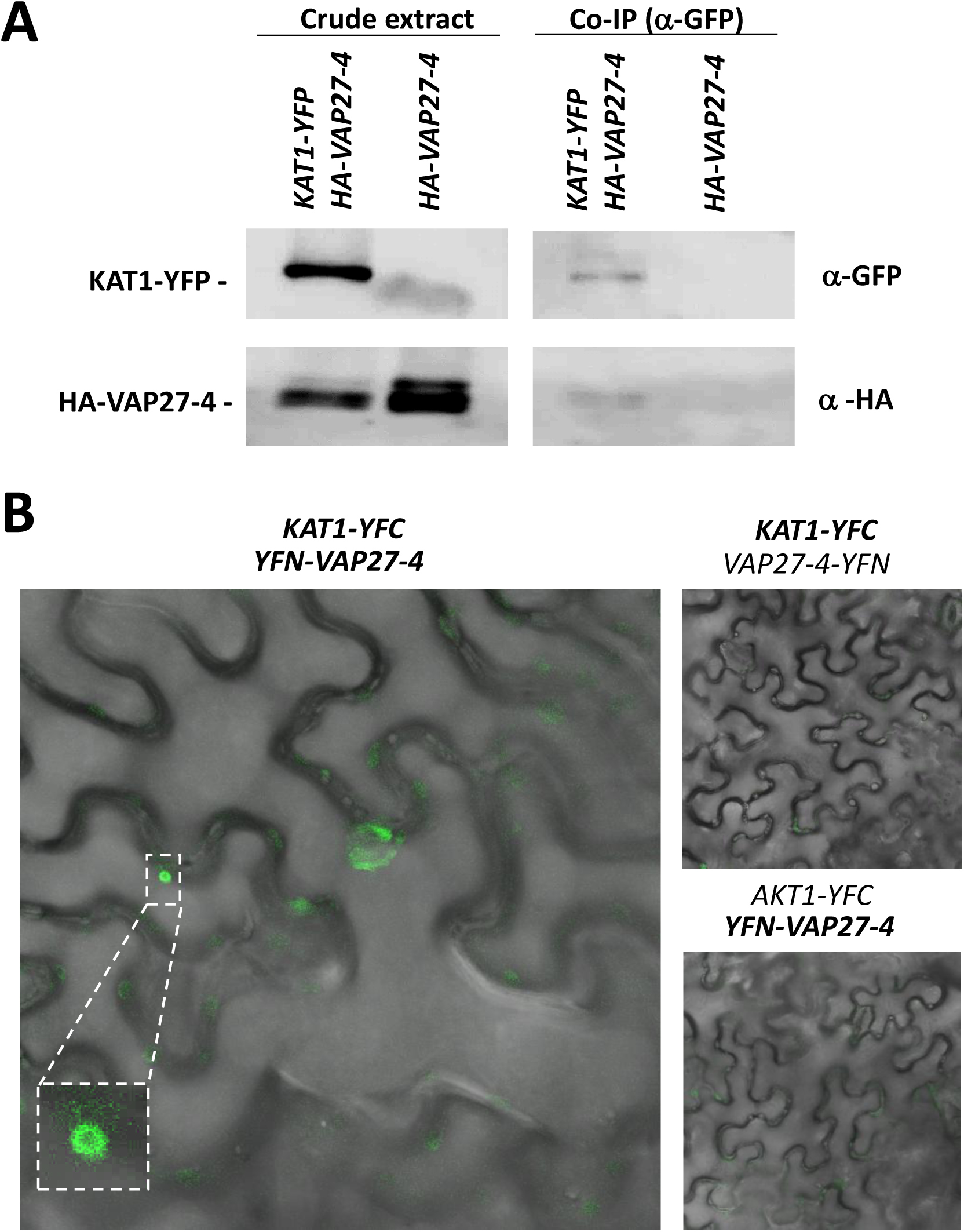
VAP27-4 and KAT1 interaction in *N. benthamiana*. **A.** Western Blot analysis of proteins co-immunoprecipitated with anti-GFP antibody from a total extract of *N. benthamiana* leaves agroinfiltrated with the *KAT1-YFP + HA-VAP27-4* or the *HA-VAP27-4* construct. The crude extract and the proteins retained in the anti-GFP beads (Co-IP) were analyzed for both constructs with anti-GFP (upper panels) and anti-HA (bottom panels) antibodies. Representative images of three independent experiments with three biological replicates are shown. The resulting bands are indicated on the left (theoretical size: KAT1-YFP, 100 kDa; HA-VAP27-4 39 kDa). **B**. Images of representative cells of *N. benthamiana* leaves agroinfiltrated with the *KAT1-YFC + YFN-VAP27-4* construct and the negative controls *KAT1-YFC + VAP27-4-YFN* and *AKT1-YFC + YFN-VAP27-*4, obtained using fluorescence confocal microscopy. Images were taken under the same conditions. Representative images of three independent experiments are shown. The merge of the YFP fluorescence in green and the bright field is shown.

Interestingly, we did not observe this interaction when YFN was fused to the C-terminus of VAP27-4. It has been reported for VAP27 family members have a C-terminal TMD domain responsible for anchoring them to the ER (Wang et al. 2016). Our data indicate that an unperturbed C-terminus of VAP27-4 is required for the KAT1 interaction.

### Dynamic localization of VAP27-4

Given the position-dependence of VAP27-4 fusions to interact with KAT1, we next analyzed the subcellular localization of YFP-VAP27-4 and VAP27-4-YFP in more detail. The C-terminal YFP fusion of VAP27-4 was described to localize to the PM, ER and EPCS as static dots (Wang et al. 2016). We confirmed that both versions co-localized with PM and ER, but not Golgi markers (Fig. S5). However, we observed different localizations for the C-terminal- and N-terminal-tagged versions of VAP27-4 (Fig. 3 and S6). The C-terminal-tagged version reproduced the static dots co-localizing with the PM, ER and EPCSs but not endosomes, whereas the N-terminal-tagged version is also observed in dot structures co-localizing with the PM, ER and both EPCSs and endosomes (Fig. 3 and S5). Composite confocal fluorescence microscopy Z-stack images reveal additional dot structures in the cytosol for the YFP-VAP27-4 fusion (Fig. S6). Interestingly, whereas the C-terminal fusion is static, the N-terminal-tagged version of VAP27-4 is observed as dot structures that actively move between the PM and the ER (Video S1).

**Figure 3.**
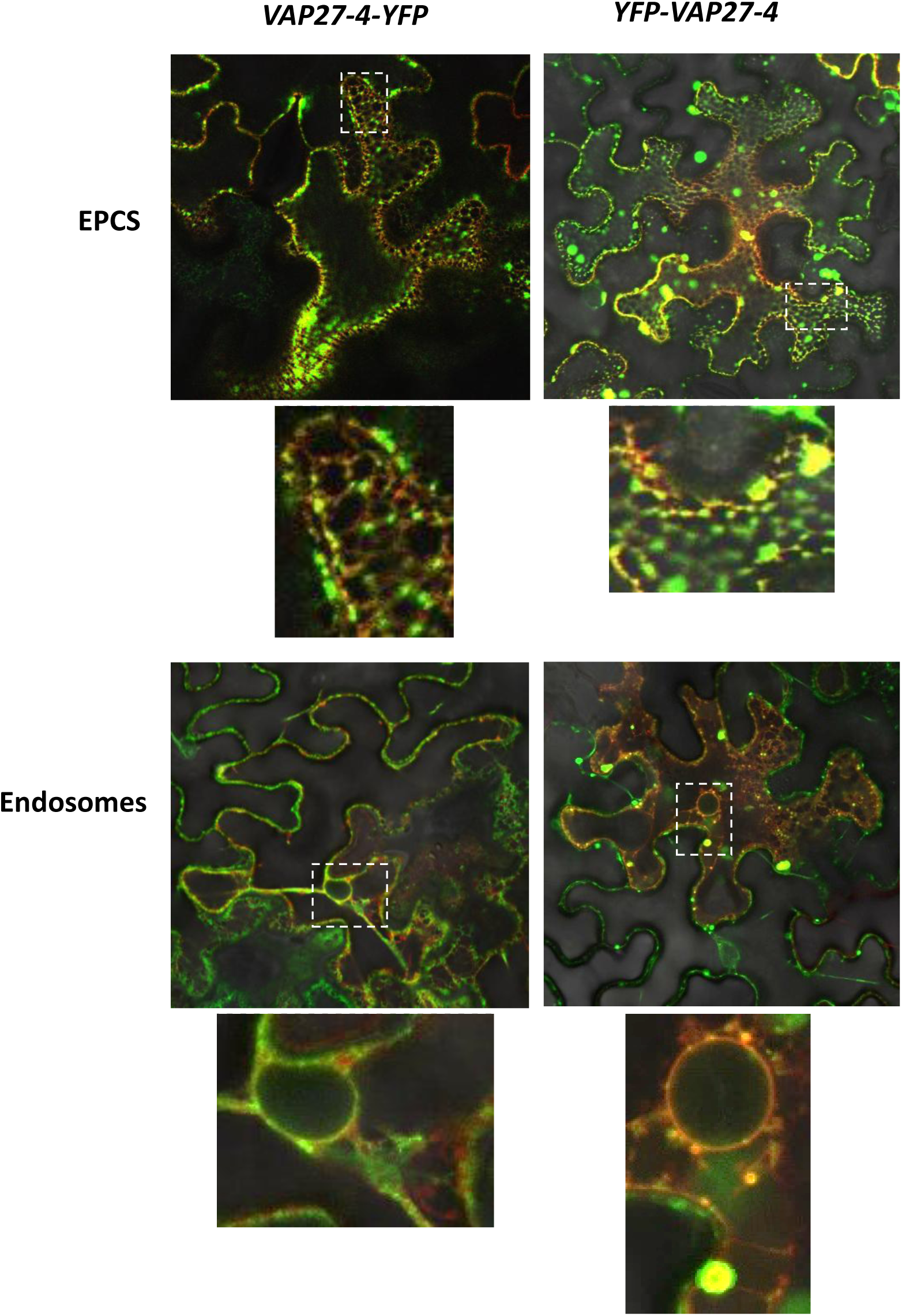
VAP27-4 subcellular co-localization with EPCSs and endosomes in *N. benthamiana*. Images of representative cells of *N. benthamiana* leaves agroinfiltrated with *VAP27-4-YFP or YFP-VAP27-4* construct together with the indicated subcellular marker (EPCS, *SYT1-RFP-T*; Endosomes, *ChFP-VAMP727*), obtained using fluorescence confocal microscopy. Representative images of two independent experiments are shown. The merge of the YFP fluorescence in green, the Cherry fluorescence in red and the bright field are shown. Zooms in the indicated areas are shown below each image.

Importantly, we confirmed these observations in Arabidopsis plants stably transformed with both tagged versions of VAP27-4. We observed subcellular localization at the PM and ER, in epidermic and guard cells (Fig. S7A) and we could also confirm the dynamic movement of the N-terminal-, but not the C-terminal-tagged version moving between the PM and the ER (Fig. S7B).

### VAP27-4 negatively regulates KAT1 in heterologous systems

Having confirmed the interaction of VAP27-4 with KAT1 (and KAT2) in planta, we next examined whether co-expression of this interactor affects KAT1 function. The first experiments were functional complementation assays in yeast. Upon co-expression of VAP27-4 and KAT1 in the *trk1 trk2* K^+^ transport-deficient mutant strain, we observed several phenotypes consistent with negative regulation of KAT1. We observed a clear decrease in growth in both solid and liquid media under limiting K^+^ conditions (5 mM solid media, 2 mM liquid media) when comparing strains co-expressing VAP27-4 or expressing KAT1 alone (Fig. 4A and 4B). We also tested the growth profile of these strains in response to toxic concentrations of LiCl. KAT1 expression in the *trk1 trk2* mutant improves K^+^ uptake and so reduces the entry of Li^+^ into the cells, leading to a tolerance phenotype (Yenush et al. 2002) . The growth rate of strains co-expressing VAP27-4 is reduced compared to those expressing KAT1 alone (Fig. 4C). BAG4 is a previously characterized KAT1 interactor that acts as a chaperone in plants increasing KAT1 PM accumulation (Locascio et al. 2019). It is used here as a specificity control.

**Figure 4.**
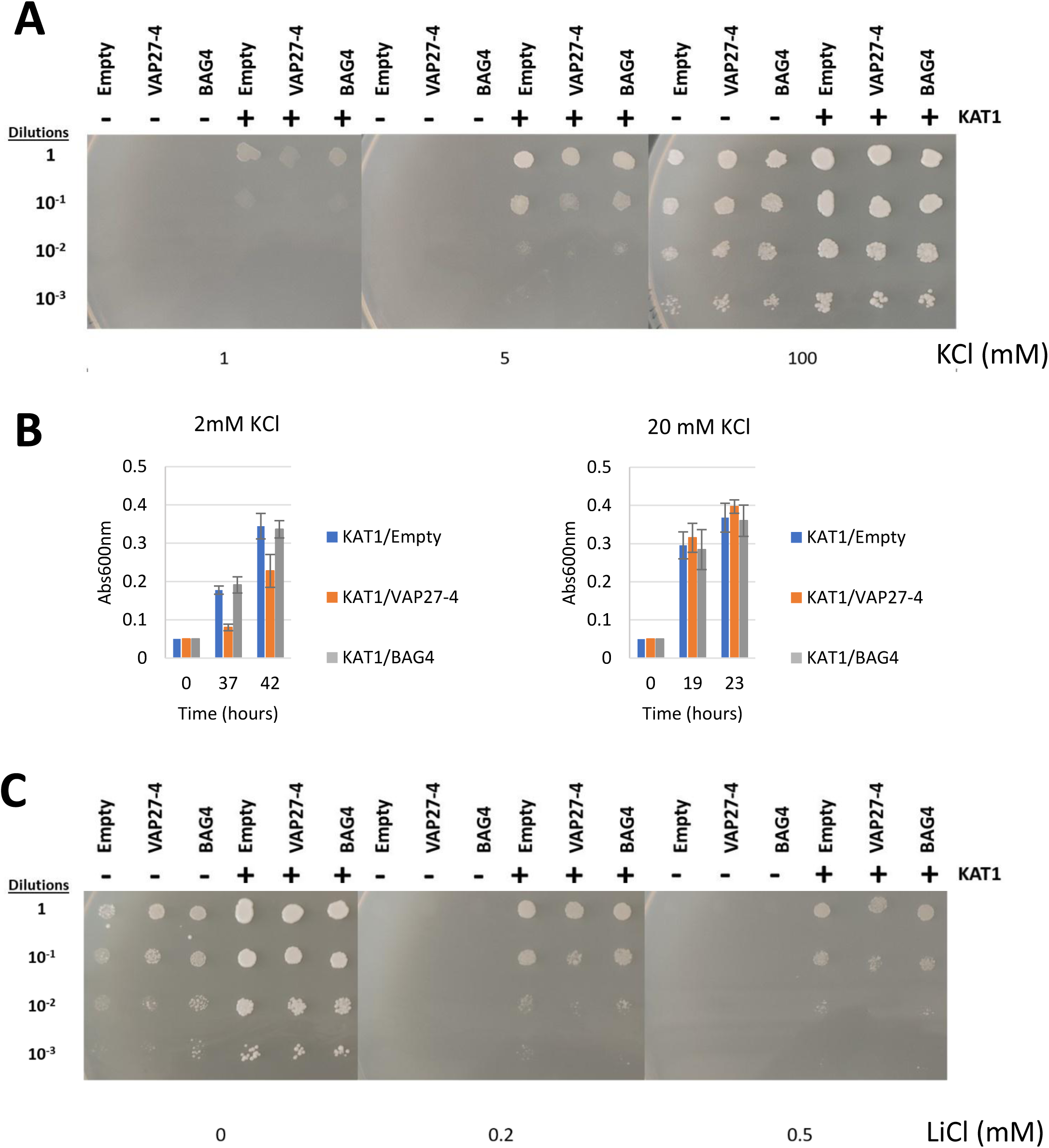
Effect of VAP27-4 on KAT1 activity in yeast. **A.** The *trk1 trk2* yeast strain was transformed with the indicated plasmids, and individual colonies were grown to saturation and serial dilutions were spotted onto low potassium minimal media supplemented with the indicated concentrations of KCl. Images were taken after 72 hours of growth at 28 °C. identical results were observed for 3 independent clones. **B.** The same strains were grown in liquid media with the indicated amount of KCl and the absorbance was measured at the indicated times. The graph shows the average value for 4 replicates and the error bars represent the standard deviation. **C**. The same strains were grown in media that was supplemented with 20 mM KCl and the indicated concentrations of LiCl. Images were taken after 72 hours of growth at 28 °C. identical results were observed for 3 independent clones.

We next tested the effect of VAP27-4 co-expression on KAT1 channel activity using two-electrode voltage clamp experiments in Xenopus oocytes. As shown in Fig. 5, we observed a decrease in KAT1 currents when VAP27-4 was co-injected at days 3, 4 and 5 post-injection. Similar results were seen for KAT2 currents upon VAP27-4 co-expression, although the effect was attenuated at day 5 in this case (Fig. S8). These electrophysiological analyses support the observations made in yeast and suggest that VAP27-4 may act as a negative regulator of KAT1 (and KAT2).

**Figure 5.**
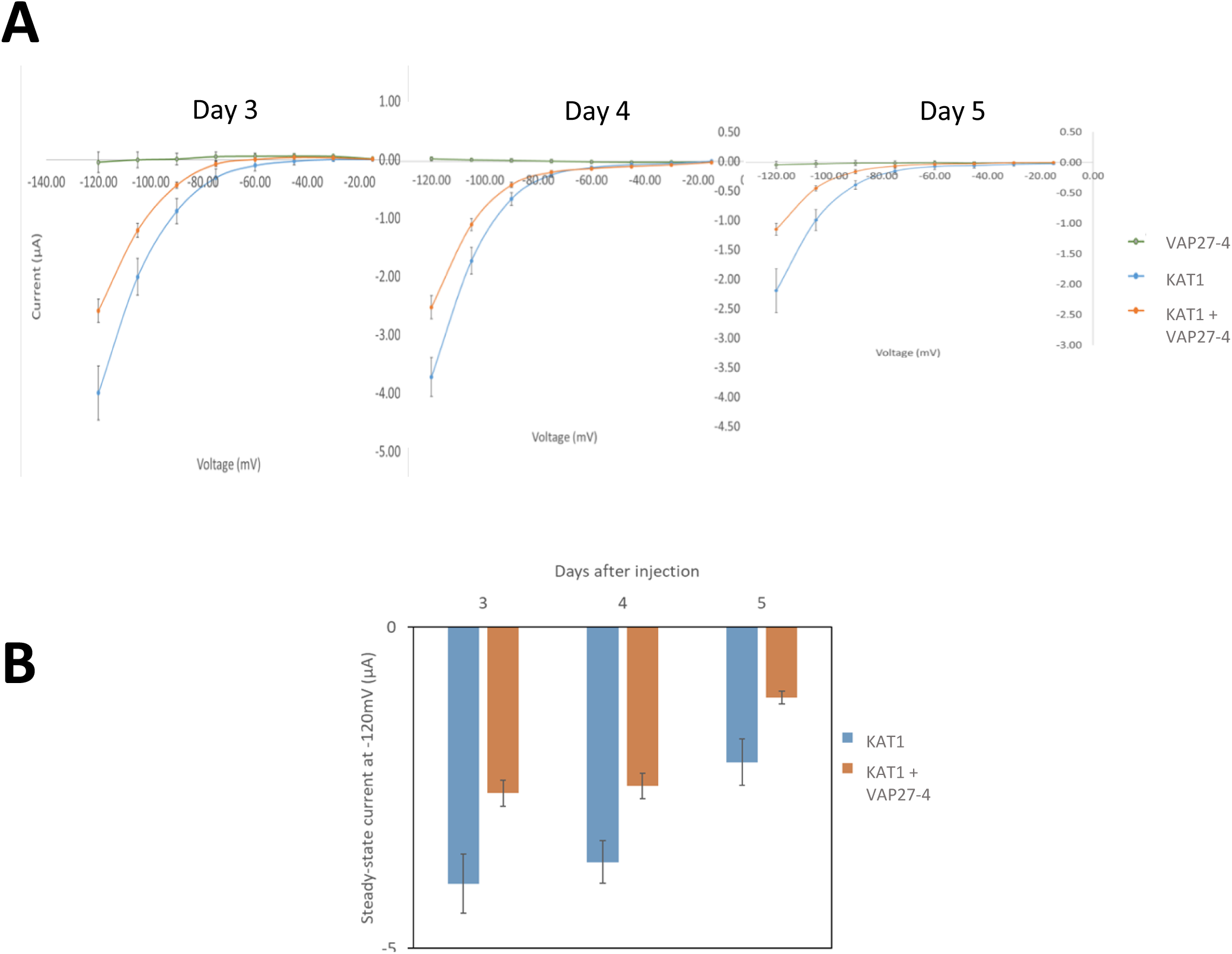
Effect of VAP27-4 on KAT1-mediated currents in Xenopus oocytes. **A.** KAT1 current (I) –voltage (V) relationships of oocytes co-expressing KAT1 and VAP27-4 (orange line), expressing KAT1 alone (blue line) or VAP27-4 alone (green line) at days 3, 4 and 5 after injection. **B.** Average steady-state current values at -120 mV for KAT1 and KAT1+VAP27-4 at 3, 4 and 5 days after injection.

### VAP27-4 participates in KAT1 endocytosis

Given the known role of VAP27 family members in endocytosis and considering the data obtained in the heterologous systems, we next investigated the possible participation of VAP27-4 in the endocytosis of the KAT1 channel. For these experiments, we generated transgenic Arabidopsis lines expressing KAT1-YFP in WT (WT*) and *vap27-4* mutant lines (KO*). WT KAT1-YFP lines were also transformed with VAP27-4 to determine the effect of overexpression (OE*). The expression levels of *VAP27-4* in these OE* lines were confirmed by RT-qPCR and immunodetection (Fig. S9A and S9C). To test the effect on KAT1 endocytosis, we performed confocal fluorescence microscopy studies on 10-day-old seedlings that were treated with 100 µM ABA and 50 µM Brefeldin A (BFA) for 1 h. As mentioned in the introduction, ABA is known to mediate KAT1 internalization (Sutter et al. 2007) and BFA has been widely used to block endosomal recycling back to the membrane, creating BFA bodies that can act as a readout for endocytosis (Lam et al. 2007). This approach has been successfully applied to study KAT1 endocytosis (Meckel et al. 2004; Sutter et al. 2007). As shown in Fig. 6, the *vap27-4* line (KO*) reduced KAT1 endocytosis, and its overexpression, in two independent lines (OE*), enhanced it. This interpretation is the same considering either the number of vesicles/stomata or the ratio of internal/PM fluorescence. No significant differences were observed for KAT1-YFP internalization in the absence of ABA treatment in the WT and *vap27-4* lines (Fig. S10) or in the total fluorescence/stomata (Fig. S11). Our results show that VAP27-4 expression positively influences KAT1 endocytosis in the guard cells of transgenic Arabidopsis plants.

**Figure 6.**
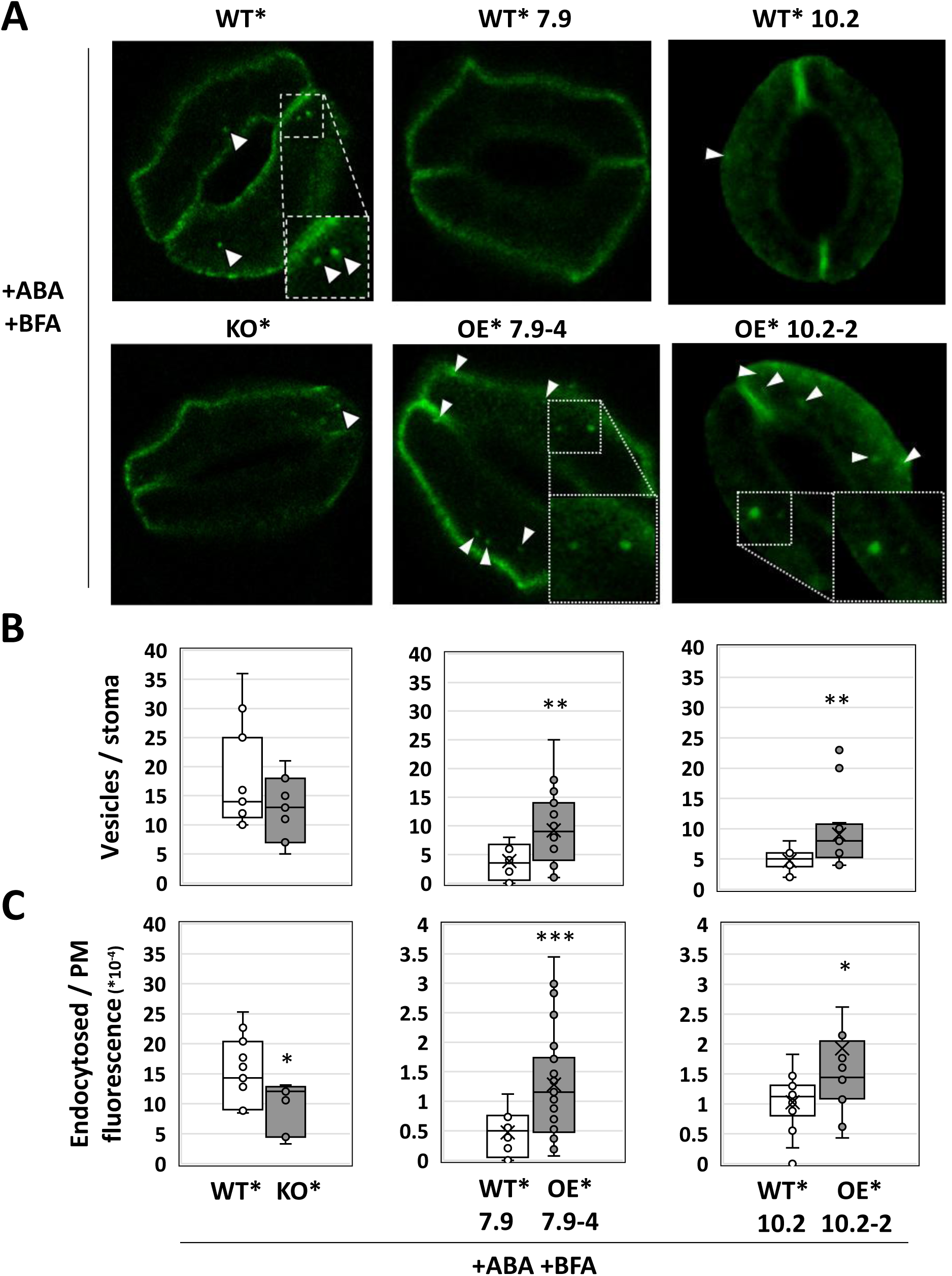
Effect of VAP27-4 on KAT1 endocytosis in Arabidopsis guard cells. **A.** Representative confocal images of a section of a stomata from leaves of *A. thaliana* WT, *vap27-4* KO and two independent *35S::HA-VAP27-4* OE lines, all of them expressing *35S::KAT1-YFP* (*). Seedlings grown for 10 long days were treated with 100 µM ABA and 50 µM BFA for 1 h. The YFP fluorescence in green is shown. White arrowheads point to KAT1-YFP fluorescence within the cell. **B.** Box-whisker plots of the number of vesicles with KAT1-YFP per stomata, in the whole stomata analyzed in panel A. **C.** Box-whisker plots of endocytosed KAT1-YFP fluorescence versus KAT1-YFP fluorescence in PM, in the whole stomata analyzed in panel A. Data were obtained from at least three different seedlings, measuring a total of seven stomata or more per line and condition. Statistical analysis of each sample with respect to WT: ***, p < 0.001; **, p < 0.01; *, p < 0.05.

### Estimating guard cell turgor pressure

To determine more directly if VAP27-4-mediated regulation of KAT1 has physiological effects on the turgor of guard cells, we used cavitation bubble manometry (CBM). This technique uses a laser to create microbubbles in guard cells, the radii of which correlate with the turgor present in the guard cells, so that, the smaller the radii, the higher the turgor pressure due to restraints on microbubble expansion (Brodersen et al. 2025). We used this approach to estimate guard cell turgor pressure in the transgenic lines. We observed an expected reduction in microbubble radius in all plants upon light exposure, indicating an increase in guard cell turgor pressure as stomata opened (Fig. 7). However, our results demonstrate that overexpression of KAT1 maintains higher guard cell turgor 1 h before light exposure compared to the WT genotype, and that genotypes co-expressing VAP27-4 with KAT1 exhibited similar changes in guard cell turgor pressure compared to the WT between all measured timepoints (Fig. 7). Microbubble radius differed significantly among measured timepoints (ANOVA p < 0.001), genotypes (ANOVA p = 0.008), with significant interactions between both factors (ANOVA p = 0.024). We also found that patterns in guard cell turgor pressure across the measured timepoints departed most considerably in the two lines overexpressing *KAT1-YFP* (WT*) from those of the WT genotype (WT) while turgor pressure patterns in lines *kat1-2* and the 2 lines co-expressing *KAT1-YFP* and *HA-VAMP27-4* (OE*) most closely resembled those of the WT genotype (Fig. 7). Across all lines excluding *WT 35S:KAT1-YFP L2 (WT*10.2)*, microbubble radii were significantly smaller at 3 h after light exposure compared to 1 h before light exposure indicating an increase in guard cell turgor pressure in response to light (Fig. S12). At 3 h after light exposure, no differences were observed in microbubble radii, and thus guard cell turgor pressure, between lines (Fig. 7). At 9 h after light exposure (i.e., 1 h after the end of the photoperiod) only the two lines overexpressing *KAT1-YFP* (WT*) showed significantly smaller microbubble radii compared to all other lines and therefore higher guard cell turgor pressure compared to the WT genotype (Fig. 7). The WT and lines *kat1-2* and the two lines co-expressing *KAT1-YFP + HA-VAP27-4* (OE*) did not differ in microbubble radius at 9 h after light exposure. Taken together, this technique indicates that co-expression of VAP27-4 mitigates the effect of KAT1 overexpression observed in guard cell turgor 1 h after the lights go off. This is consistent with VAP27-4 increasing KAT1 endocytosis, reducing K^+^ uptake and thus reducing guard cell turgor.

**Figure 7.**
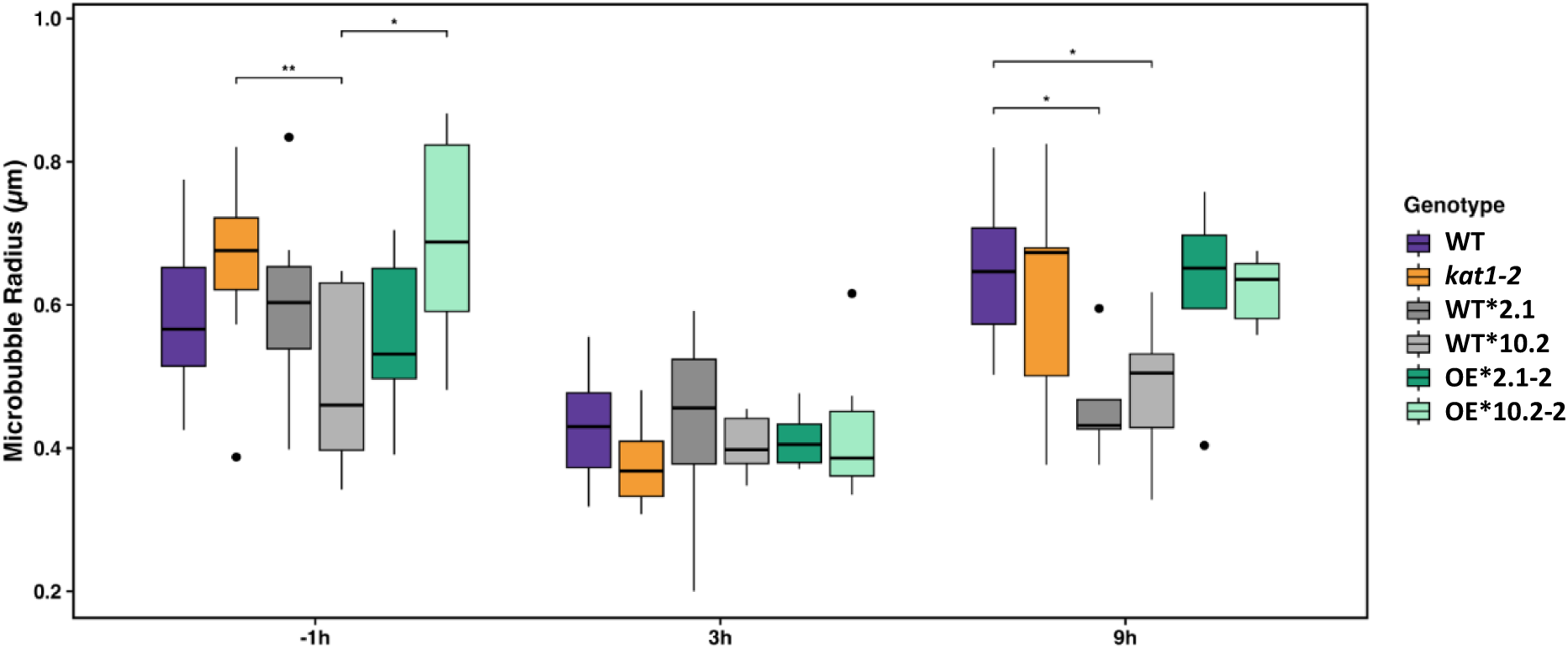
Changes in guard cell turgor in response to light. Guard cell microbubble radius across genotypes for WT, *kat1-2*, two independent WT *35S::KAT1-YFP* lines (WT*2.1 and WT*10.2), two independent WT *35S::KAT1-YFP* + *35S::HA-VAP27-4* (OE*2.1-2 and OE*10.2-2), and measured timepoints including -1h prior to light exposure, 3h following light exposure, and 9h following light exposure (i.e., one hour following the end of the photoperiod). Smaller microbubble radii indicate higher guard cell turgor pressure. Boxes and whiskers show the median, interquartile ranges, and 1.5x the interquartile range. Significant Tukey-adjusted pairwise comparisons between genotypes within each timepoint are indicated with brackets and asterisks. *p < 0.05; **p < 0.01.

### VAP27-4 participates in stomata closure

Finally, to determine if loss-or gain-of-function of *VAP27-4* has effects on the whole plant level, we measured stomatal dynamics in response to light in mutant and over-expression lines, as described in Materials and Methods. A representative experiment is shown in Fig. 8A, depicting the distribution of stomatal aperture measurements for each time and line, and Fig. 8B shows a graph representing the average values for three different experiments. We observed that the loss-of-function of *VAP27-4* enhanced the stomata opening at 3 and 6 h of light treatment. On the other hand, the corresponding complemented line overexpressing *VAP27-4* exhibited slightly faster closure at 6 h. These results suggest that VAP27-4 plays an important role in stomatal dynamics in response to light.

**Figure 8.**
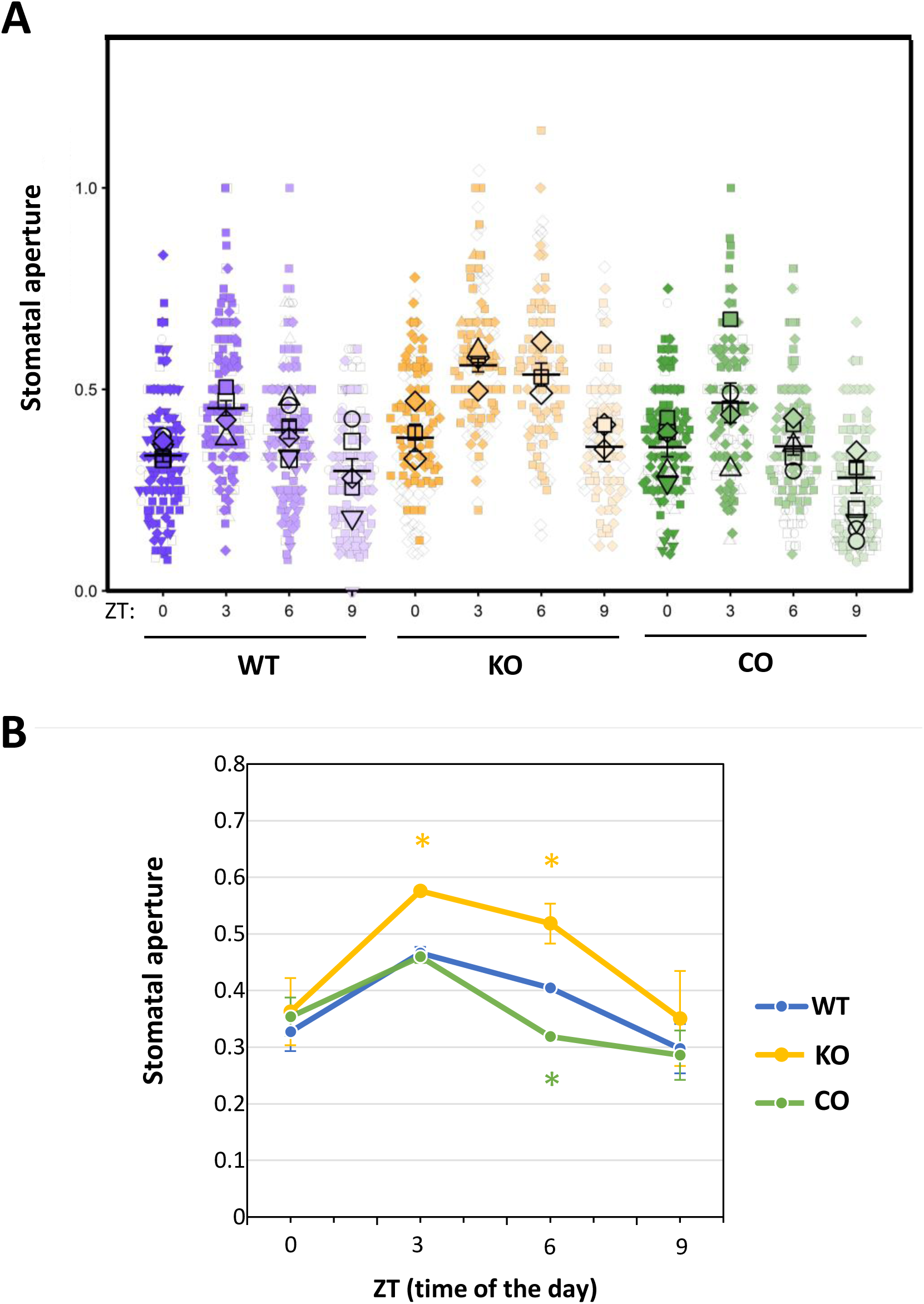
Effect of VAP27-4 on stomatal opening in Arabidopsis. Stomatal aperture (width/length) in cotyledons of 4-day-old *A. thaliana* WT*, vap27-4 (*KO) and *vap27-4* + *35S::HA-VAP27-4 (CO)* line. Seedlings were grown under short-day conditions, and photos for measurements were taken 0, 3, 6 and 9 h after lights switch on. The means and the standard deviations of at least 200 stomata, from at least three seedlings of 3 different experiments, are shown in panel **A**. The means and the standard deviations of the three independent experiments are shown in panel **B**. Statistical analysis of each sample with respect to the WT of the same condition is shown in panel B: *, p-value ≤ 0.05.

## Discussion

In the present study we present evidence showing that the VAP (<u>V</u>AMP (vesicle-associated membrane proteins) <u>A</u>ssociated <u>P</u>rotein) family member, VAP27-4 mediates the endocytosis of KAT1 K^+^ channel through a direct physical interaction (Fig. 9). This interaction was initially identified in a high-throughput screen in yeast using KAT1 as the bait protein. We previously reported our results characterizing three other interactors from this screening, RPT2, MEE31 and BAG4, which are all positive regulators of KAT1 favoring channel accumulation at the PM (Locascio et al. 2019, 2025; González-García et al. 2024). In addition to these proteins with chaperone-like functions, KAT1 is known to be regulated by several other proteins, most notably SYP121 and VAMP721, which are involved in KAT1 exocytosis and recycling back to the PM and, in the case of SYP121, directly favoring channel opening (Eisenach et al. 2012; Zhang et al. 2015, 2017; Lefoulon et al. 2018). Here, we show that VAP27-4 acts as a negative regulator of KAT1 in yeast and oocytes. It is conceivable, based on our results in these heterologous systems, that VAP27-4 may exert a direct effect on KAT1 gating, especially if we consider that the molecular machinery that would be required for this endocytic regulation is not highly conserved. Since SYP121 is involved in exocytosis and positively affects gating, it is tempting to speculate that VAP27-4 may have a similar dual role facilitating endocytosis and negatively affecting channel activity.

**Figure 9.**
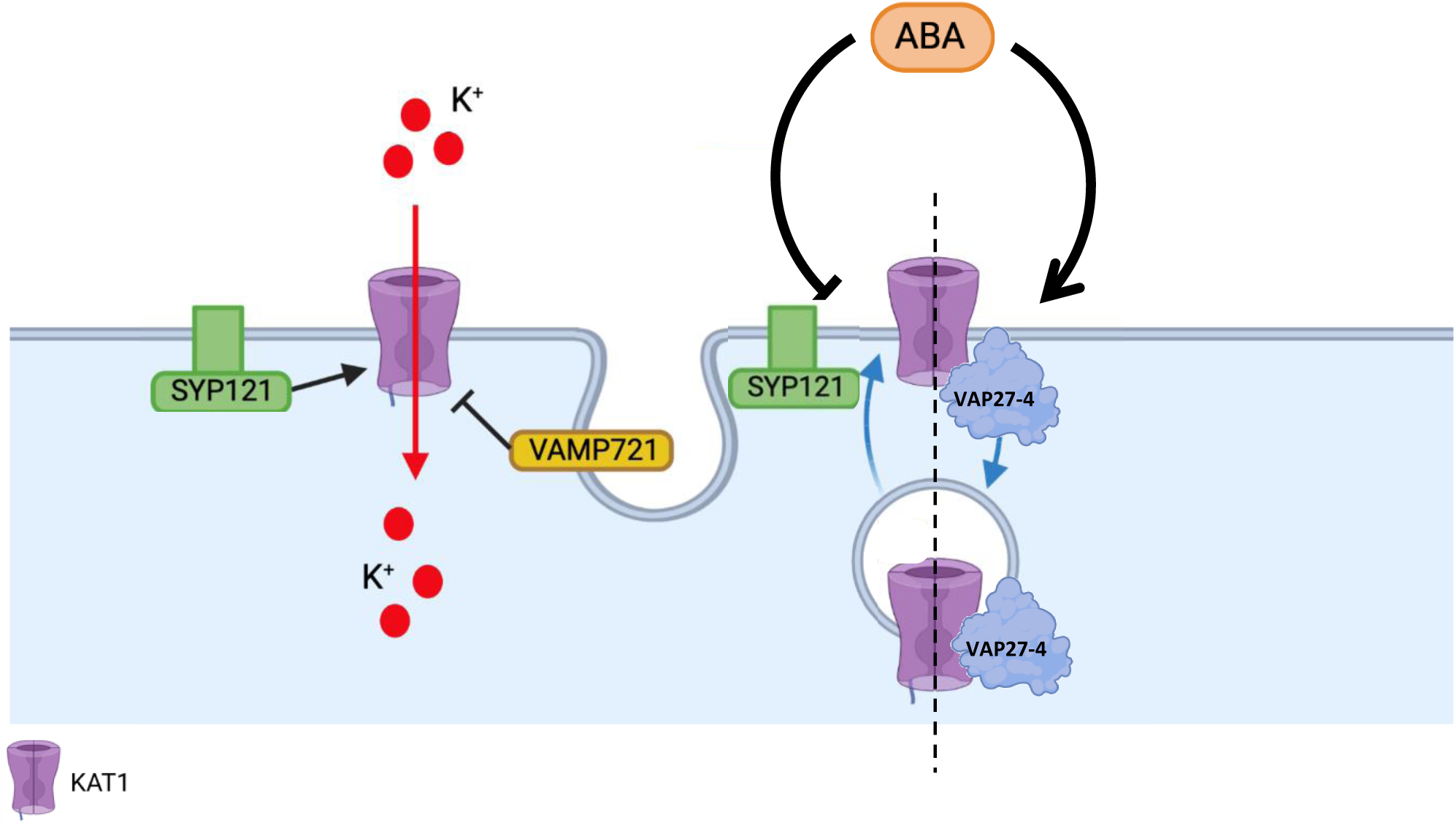
Working model for VAP27-4-mediated KAT1 regulation. ABA-dependent endocytosis of KAT1 is mediated by VAP274, whereas SYP121 and VAMP721 play a role in KAT1 accumulation and recycling to the plasma membrane (Sutter et al. 2007; Eisenach et al., 2012; Zhang et al., 2015; Lefoulon et al., 2018.).

Our results indicate that fusing proteins to the C-terminus of VAP27-4 impedes its interaction with KAT1. Among the conserved structural components of the VAP27 family is the C-terminal TMD required for ER association (Wang et al. 2016; Stefano et al. 2018). We show here contrasting behaviors for both the subcellular localization and the movement of VAP27-4 with YFP fusions in the N-or C-termini. It appears that, in addition to inhibiting the interaction with KAT1, an unperturbed C-terminal TMD is necessary for the localization of this protein in vesicles that shuttle between the ER and the PM (Video S1). The C-terminal fusion of VAP27-4 is still associated with the ER, PM and EPCSs, but is static. In the BiFC assays studying the physical interaction between VAP27-4 and KAT1, we do not observe this movement, suggesting that it is possible that the pool of VAP27-4 that is not bound to cargo proteins is mobile. It will be interesting to study whether this same phenomenon occurs with other VAP27 family members. Our preliminary data suggests that VAP27-1 and VAP27-3 show a similar behavior. How this dynamic localization participates in the function of these trafficking proteins warrants future studies.

The more important challenge for characterizing a new protein-protein interaction is to prove that it is physiologically relevant. According to public databases, VAP27-4 gene expression is higher and opposite to KAT1 at night (http://viridiplantae.ibvf.csic.es/circadiaNet/genes/atha/AT5G47180.html), in response to ABA in stomata (https://bar.utoronto.ca/eplant/), and in response to drought, especially late in the day (https://bar.utoronto.ca/eplant/). In other words, under the conditions mentioned, as *VAP27-4* gene expression increases, *KAT1* transcription begins to decline. Of course, these expression patterns do not demonstrate a causal relationship, but they are consistent with VAP27-4 being a negative regulator of KAT1 in guard cells and in response to different stimuli. To generate data to refute or support this hypothesis, we studied the effect of VAP27-4 loss-of-function and gain-of-function on KAT1 endocytosis in the guard cells of transgenic lines in response to ABA and in the presence of BFA. Because BFA inhibits recycling without stopping initial uptake, the rate at which cargo accumulates in these aggregates serves as a standard quantitative readout for endocytosis. We observed results consistent with VAP27-4 participating in the endocytosis of KAT1 (Fig. 9). The *vap27-4* mutant line showed reduced ABA-stimulated KAT1-YFP endocytosis, whereas two independent transgenic lines co-expressing KAT1 and VAP27-4 accumulated more KAT1-YFP internally upon phytohormone treatment.

Even with the expected changes in the VAP27-4-dependent internal accumulation of KAT1-YFP in the endocytosis assays, it could be argued that sufficient KAT1 channel activity would remain on the cell surface of the guard cells, rendering this regulatory mechanism ineffective and/or artefactual. The direct measurement of guard cell electrophysiology in Arabidopsis is highly complex. In the present study, we took an alternative approach to estimate guard cell turgor in our transgenic lines. Cavitation bubble manometry (CBM) allows for generation and measurement *in situ* of the radii of microbubbles formed as a proxy for guard cell turgor pressure (Brodersen et al. 2025). In this technique, the growth and dissolution of nucleated microbubbles created using modified laser ablation components inside the guard cells of intact plant tissue are recorded using a high-speed camera. It has been demonstrated that higher pressure reduces the maximum microbubble radius because the surrounding fluid restricts expansion, following a modified Epstein–Plesset equation. Our previous reports have shown that overexpression of KAT1 in Arabidopsis leads to increased stomatal aperture and delayed stomatal closure in response to light (Locascio et al. 2019, 2025; González-García et al. 2024; Rovira et al. 2024). Using CBM, we observed a significant decrease in microbubble radii, compared to the WT, in both transgenic lines overexpressing KAT1 at 9 hours, which is 1 hour after the lights shut off and the stomata should be closing. Importantly, this phenotype was reverted in two independent transgenic lines co-expressing VAP27-4. This data is consistent with VAP27-4 mediating increased KAT1 endocytosis, with the corresponding reduction in K^+^ influx, leading to this decrease in estimated turgor. These results provide strong evidence that VAP27-4 is indeed affecting KAT1 function in guard cells in intact plant tissue.

Finally, we were able to corroborate these observations by measuring stomatal dynamics of the loss- and gain-of-function *VAP27-4* lines. We observed a marked effect on stomatal opening in the loss-of-function mutant, showing increase aperture at both 3 and 6 hours of light, as compared with the WT. These results would be consistent with an overaccumulation of endogenous KAT1 (and possibly KAT2) at the plasma membrane of guard cells leading to an increase in K^+^ influx in response to the light-induced hyperpolarization of the membrane. On the other hand, the overexpression line showed faster closure at 6 hours. In the overexpression line, increased levels of VAP27-4 would facilitate the endocytosis of KAT1 (and possibly KAT2) towards the end of the light cycle to accelerate stomatal closure. It is important to point out that these lines do not overexpress KAT1 and so are not directly comparable to the CBM results. However, both experiments provide complementary data supporting a role for VAP27-4 in KAT1 endocytosis that is physiologically relevant.

Several outstanding questions merit further investigation. For example, our data establish a possible paradigm that could be extended to other ion transporters, for example SLAC1 and SLAH3, which are known to physically interact with KAT1 (Zhang et al. 2016). Whether SYP, VAMP or VAP27 family members are involved in their trafficking is an interesting possibility. Finally, in mammals, VAP proteins have been shown to interact with the Voltage-gated K^+^ channel Kv2 (Johnson et al. 2018). This Kv2-VAP interaction mediates the association with ER-PM contact sites (EPCS) responsible for Kv2 clustering. The phosphorylation of serine residues in the Proximal Restriction & Clustering (PRC) domain of Kv2 produces negative charges to enable VAP binding and control the clustering of Kv2 channels, which is a prerequisite for Kv2 channel-mediated ER-PM connections. Although KAT1 does not possess a recognized PRC domain, it has been reported to be phosphorylated on Thr_306_ by the SnRK2.6/OST1 protein kinase (Sato et al. 2009). This kinase is a known component of the core ABA signaling pathway. KAT1 channel activity has also been described to be regulated by phosphorylation by other kinases, especially of the CDPK family. It would be interesting to know if these phosphorylation events are important for VAP27-4 interaction and the internalization of KAT1.

## Conclusions

Our results identify the trafficking protein, VAP27-4 as a novel regulator of the endocytosis of the inward-rectifying K^+^ channel KAT1. We have demonstrated the physical and functional interaction in heterologous systems, the physiological relevance of this regulatory interaction in guard cell turgor, and a role for VAP27-4 in stomatal dynamics.

## Author contributions-CRediT statement

Conceptualization and Funding acquisition: LY and JMM; Investigation: NAC, MNC, JZ, ERC, VG, IM, LGR, JA, LC, LT, PC, EMS and LY; Data curation and figure preparation: NAC, MNC, JZ, CB, ERC, JR, LY; Writing - original draft: NAC, JZ, LY; Writing-review and editing: LY, MNC, JZ, JR, BBP, CB and JMM.

## Funding Sources

This work was supported by PAID-06-18 (NAC, JMM, LY), PID2022-137193NB-I00 (NAC, JMM, LY), PID2025-172444OB-I00 (MNC).

## Declaration of competing interest

The authors declare that they have no known competing financial interests or personal relationships that could have appeared to influence the work reported in this paper.

## Acknowledgements

Maria Pilar López and Marisol Gascón are acknowledged for excellent technical assistance. The authors would also like to thank Dr. María Jesús Marcote, Dr. Noemi Ruíz-López and Dr. Vicente Pallás for providing helpful reagents. We would like to acknowledge the Protein Expression in Xenopus Oocytes Service of the Centro de Edafología y Biología Aplicada del Segura (CEBAS – CSIC).

---

**Supplementary Table 1.**
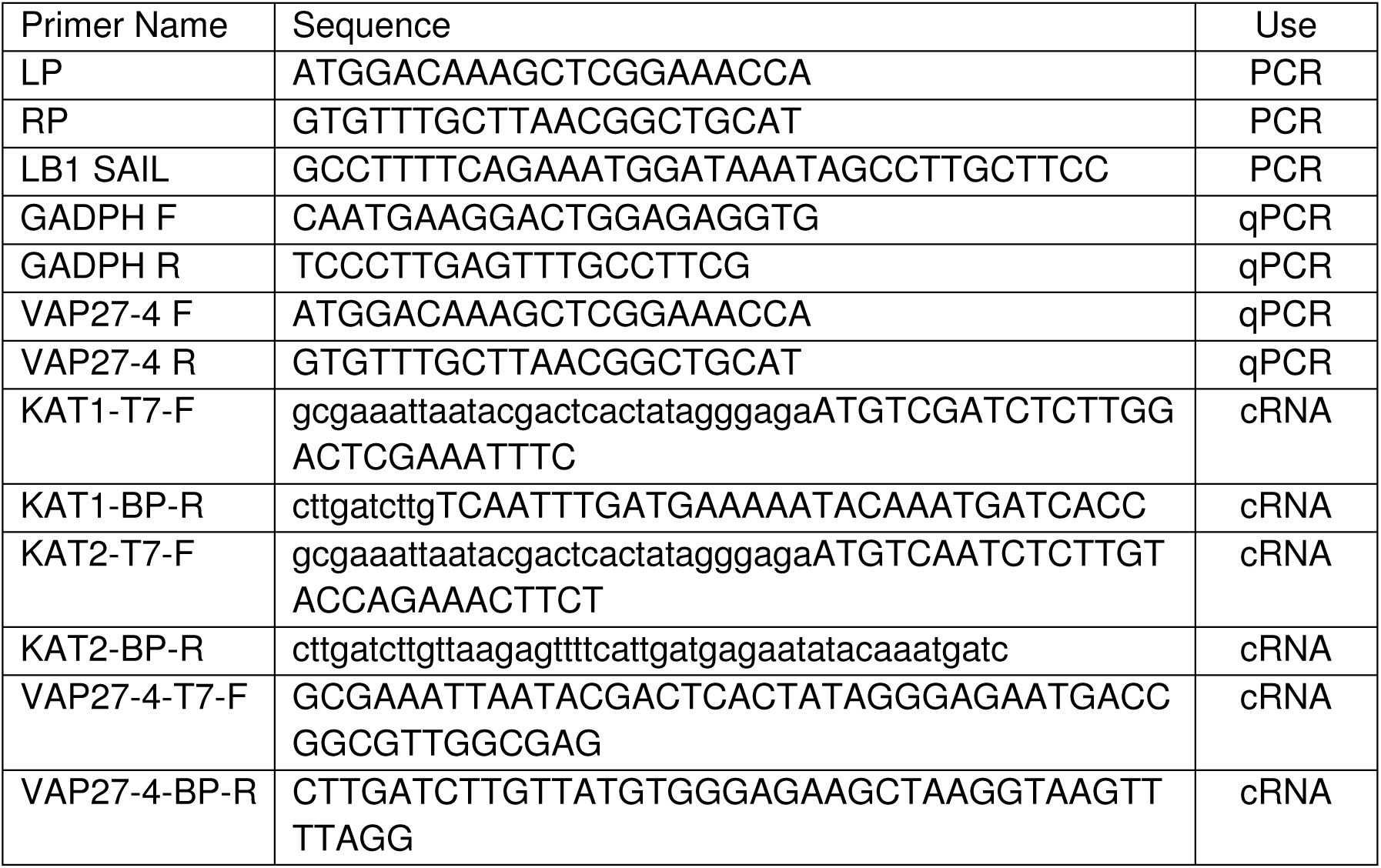
Primers used in this study.

## Supplemental Figures

**Figure S1.**
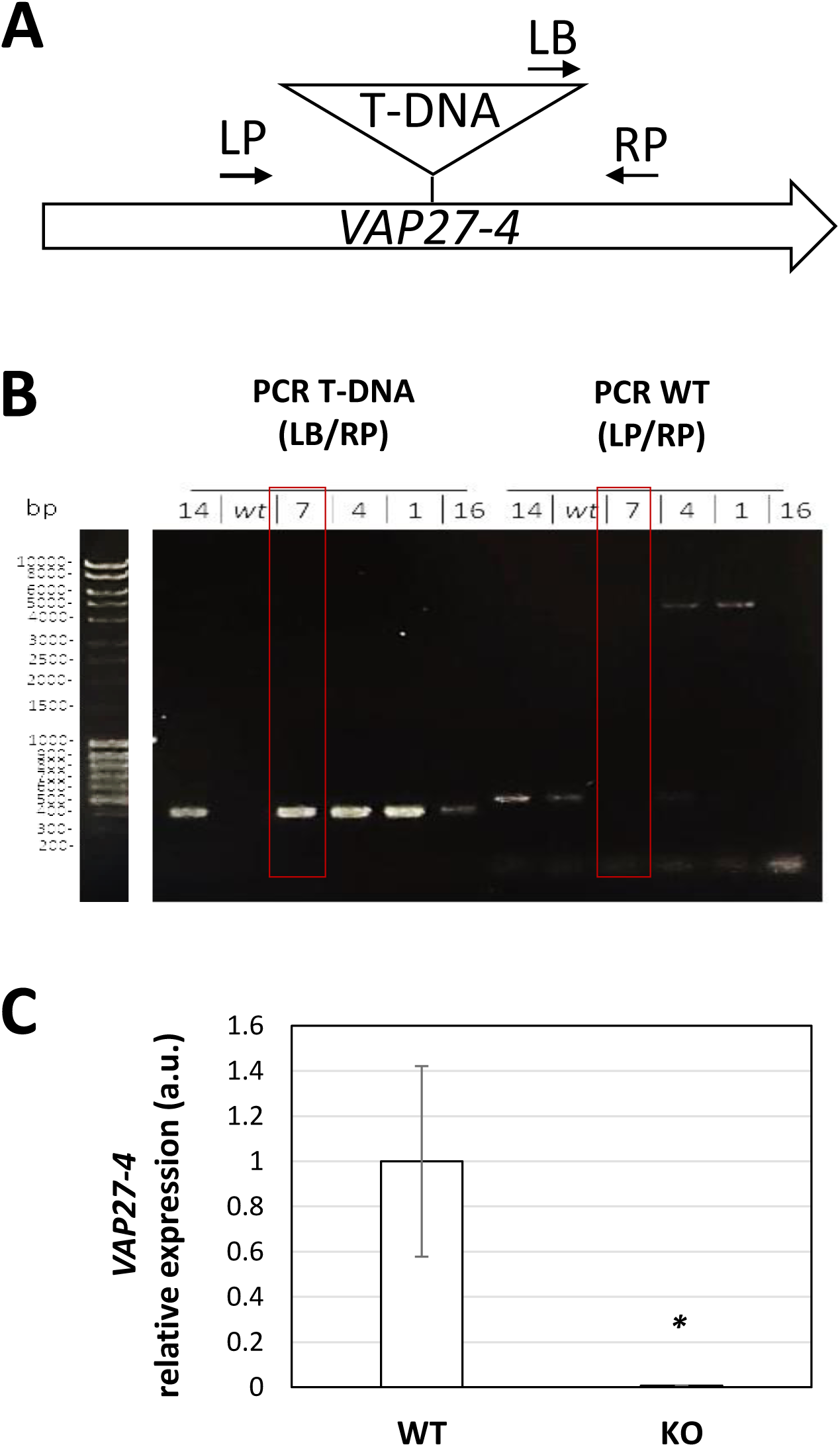
Genotyping and expression analysis of *VAP27-4* in the knockout line. **A.** Schematic diagram of the primary structure of *VAP27-4* and T-DNA insertion site. The position and name of the primers used for genotyping are indicated with continuous arrows. **B.** Genotyping of the *vap27-4* KO line. Electrophoresis of PCR products of gDNA samples from leaves of the WT and *vap27-4* KO lines analyzed by standard PCR with specific primers, as indicated. PCR T-DNA: LB1 SAIL + RP SAIL, 440 pb; PCR WT: LP SAIL + RP SAIL, 505 pb. **C.** *VAP27-4* expression levels in *vap27-4* KO line. Total RNA from leaves of adult *A. thaliana* plants grown under long-day conditions was analyzed by RT-qPCR, from WT and *vap27-4* KO lines. Expression values of the *VAP27-4* gene were normalized with respect to the *GADPH* control gene. The WT line was taken as reference (value = 1). The mean of at least two biological replicates is shown with their respective standard deviation. Statistical analysis with respect to WT : *, p-value ≤ 0.05. a.u., arbitrary units.

**Figure S2.**
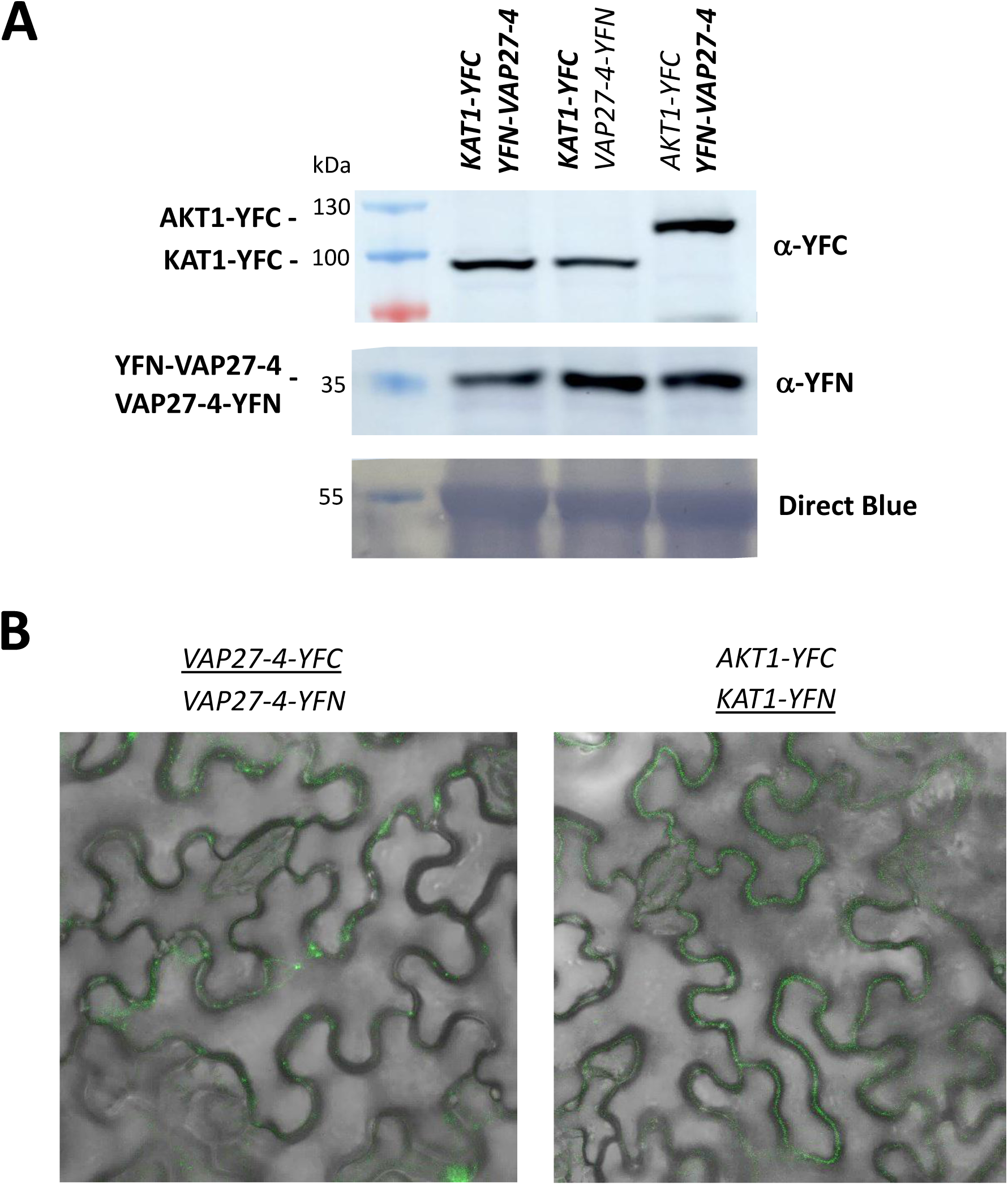
Controls of VAP27-4 and KAT1 interaction in *N. benthamiana* by BiFC assay. **A.** Western Blot analysis from total extracts of the *N. benthamiana* leaves shown in Figure 2 . The anti-YFN/YFC antibodies were used for protein detection. Direct blue staining of the membrane before antibody incubation is shown in the bottom part as a loading control. The resulting bands are indicated on the left (theoretical size: AKT1-YFC, 107.1 kDa; KAT1-YFC, 88 kDa; YFN-VAP27-4/VAP27-4-YFN, 43.67 kDa). **B.** Images of representative cells of the *N. benthamiana* leaves agroinfiltrated with the *VAP27-4-YFC + VAP27-4-YFN* or *AKT1-YFC + KAT1-YFN* construct, obtained using fluorescence confocal microscopy. Images were taken under the same conditions as in Figure 2. Representative images of three independent experiments are shown. The merge of the YFP fluorescence in green and the bright field are shown.

**Figure S3.**
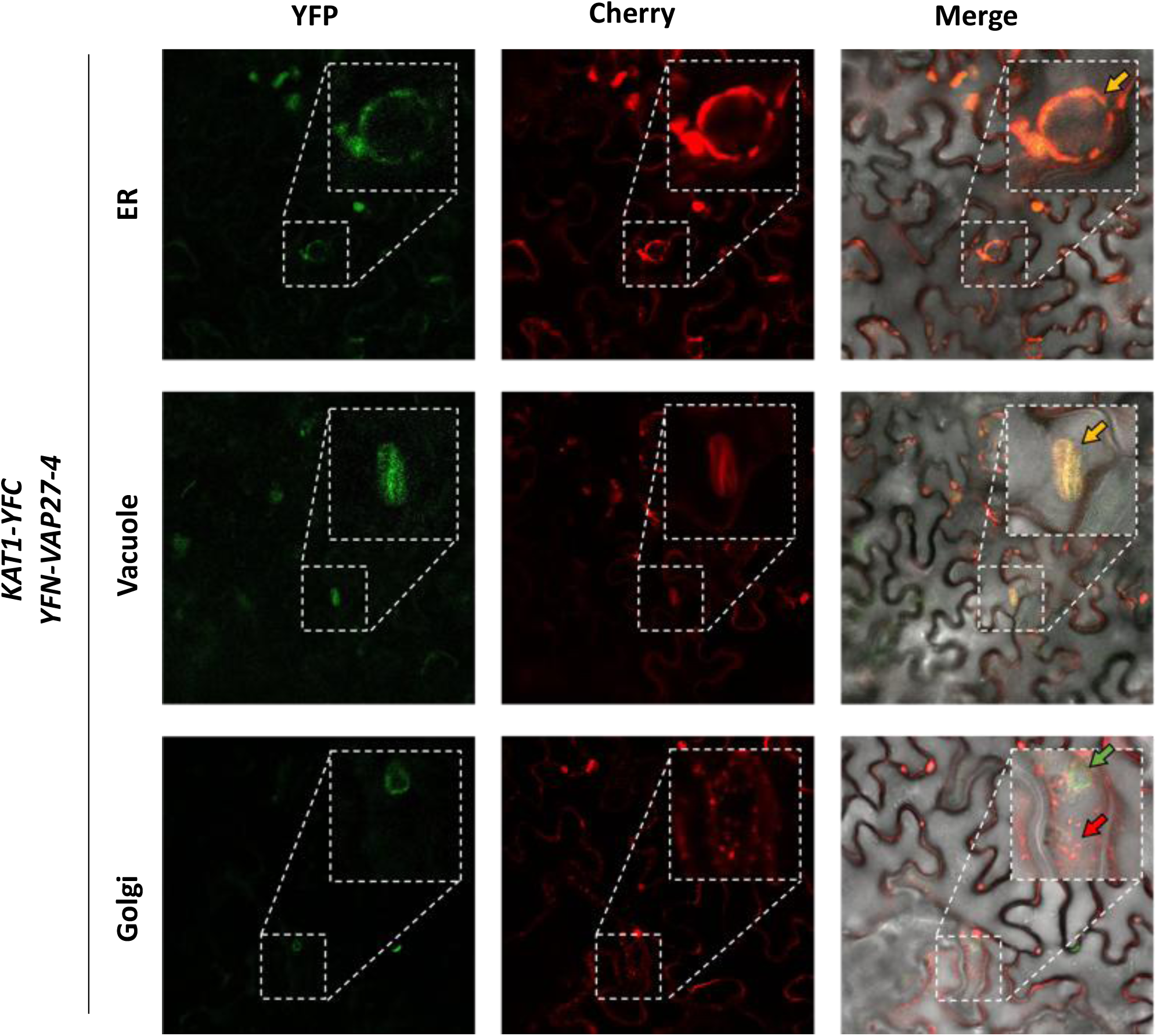
Co-localization of VAP27-4 and KAT1 interaction in *N. benthamiana*. Images of representative cells of *N. benthamiana* leaves agroinfiltrated with the *KAT1-YFC + YFN-VAP27-4* construct, together with the indicated subcellular marker (ER, *HDEL-ChFP;* vacuole, *ChFP-VAMP713;* Golgi, *pCBNoXbaSTtmdCherry*), obtained using fluorescence confocal microscopy. Representative images of three independent experiments are shown. The YFP fluorescence in green, the Cherry fluorescence in red and the merge with the bright field are shown. Yellow arrows, interaction; red and green arrows, no interaction.

**Figure S4.**
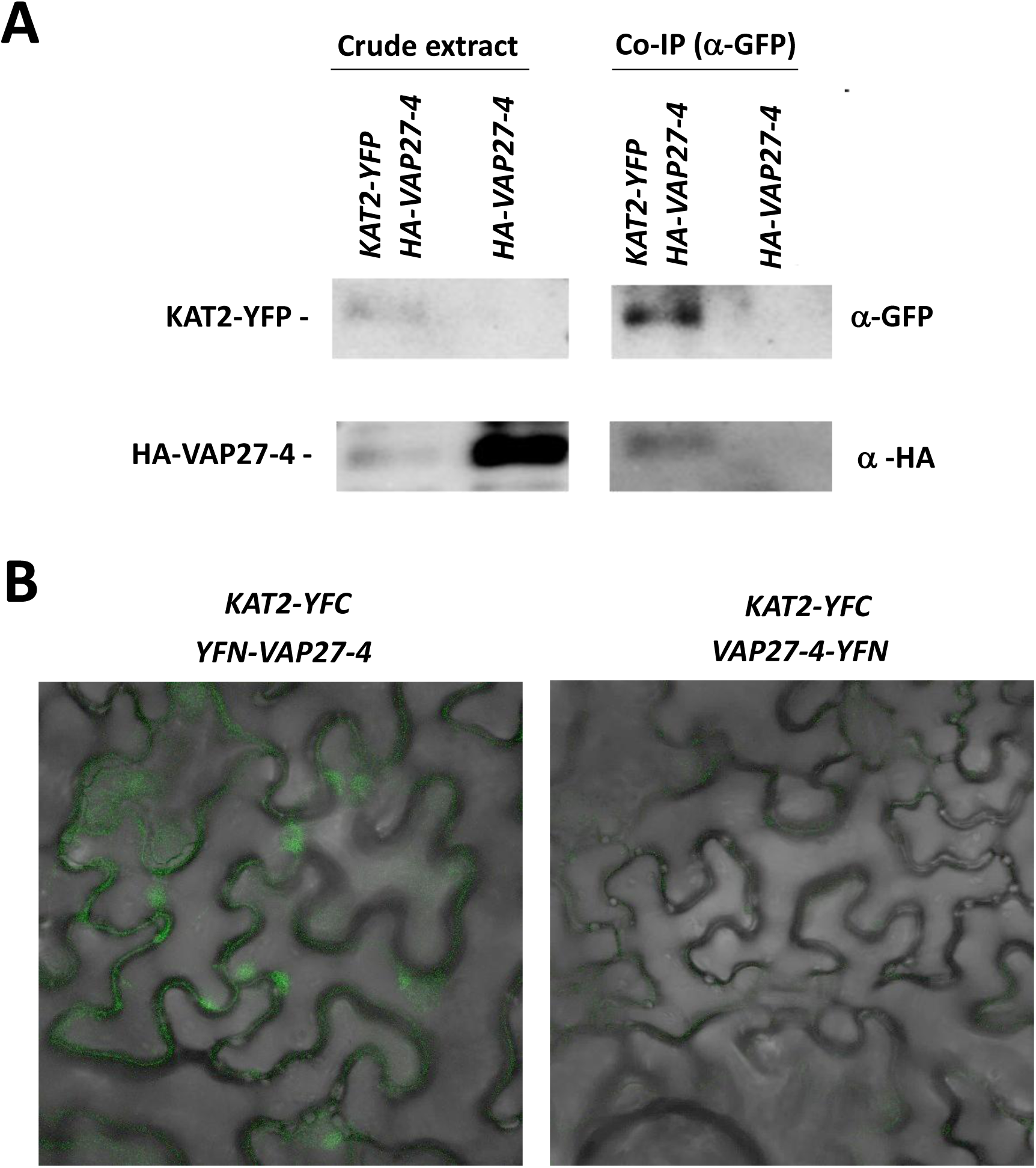
VAP27-4 and KAT2 interaction in *N. benthamiana*. **A.** Western Blot analysis of proteins co-immunoprecipitated with anti-GFP antibody from a total extract of *N. benthamiana* leaves agroinfiltrated with the *KAT2-YFP + HA-VAP27-4* or the *HA-VAP27-4* construct. The crude extract and the proteins retained in the anti-GFP beads (Co-IP) were analyzed for both constructs with anti-GFP (upper panels) and anti-HA (middle panels) antibodies. Representative images of three independent experiments with three biological replicates are shown. The resulting bands are indicated on the left (theoretical size: KAT2-YFP, 100 kDa; HA-VAP27-4 39 kDa). **B.** Images of representative cells of the *N. benthamiana* leaves agroinfiltrated with the *KAT2-YFP* construct, together with *YFN-VAP27-4 or VAP27-4* construct, obtained using fluorescence confocal microscopy. Representative images of three independent experiments are shown. The merge of the YFP fluorescence in green and the bright field are shown.

**Figure S5.**
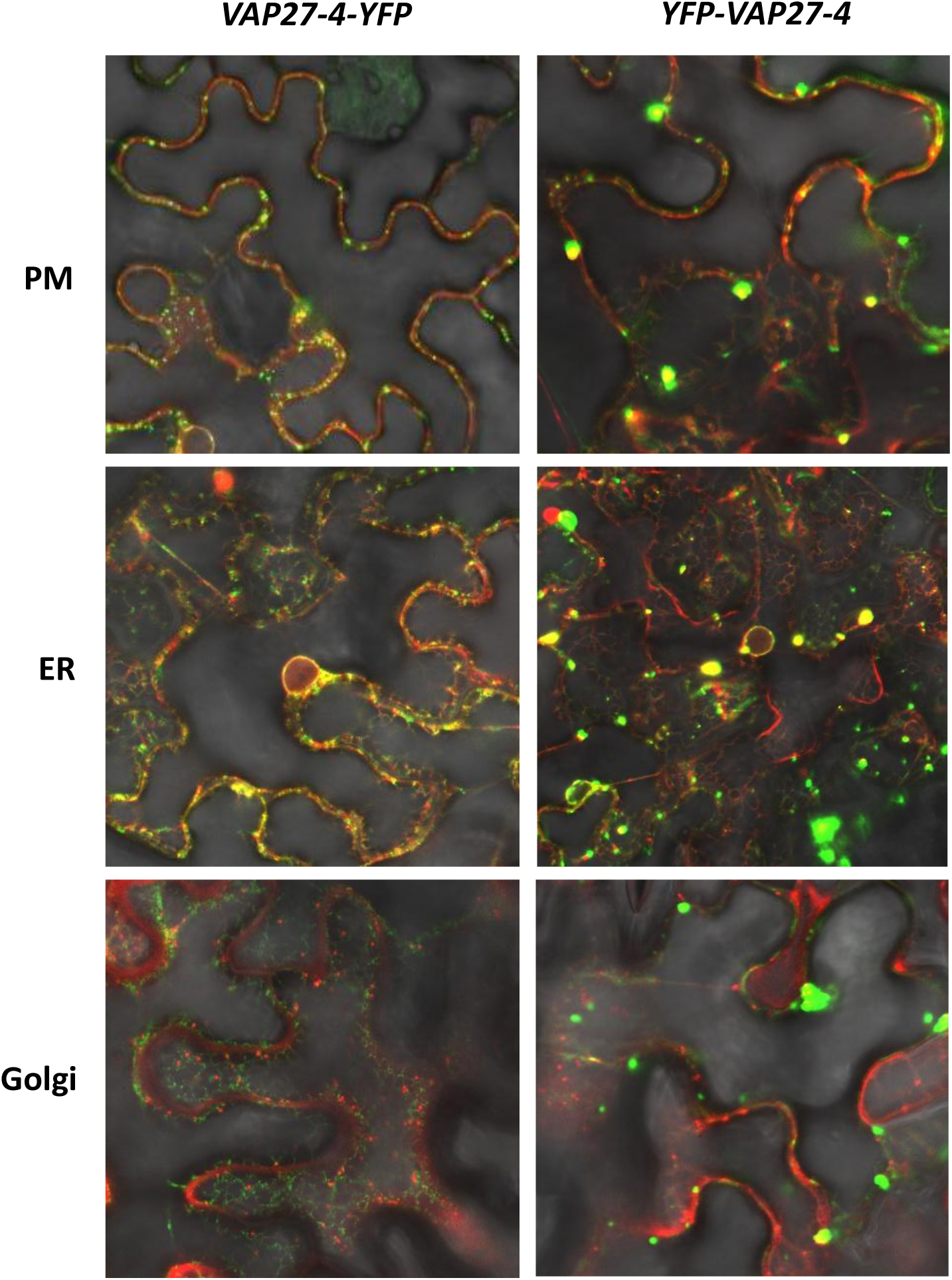
VAP27-4 subcellular co-localization with PM and ER in *N. benthamiana*. Images of representative cells of *N. benthamiana* leaves agroinfiltrated with *VAP27-4-YFP or YFP-VAP27-4* construct, together with the indicated subcellular marker (PM, *AtPIP2A-Cherry;* ER, *HDEL-ChFP;* Golgi, *pCBNoXbaSTtmdCherry*), obtained using fluorescence confocal microscopy. Representative images of three independent experiments are shown. The YFP fluorescence in green, the Cherry fluorescence in red and the merge with the bright field are shown.

**Figure S6.**
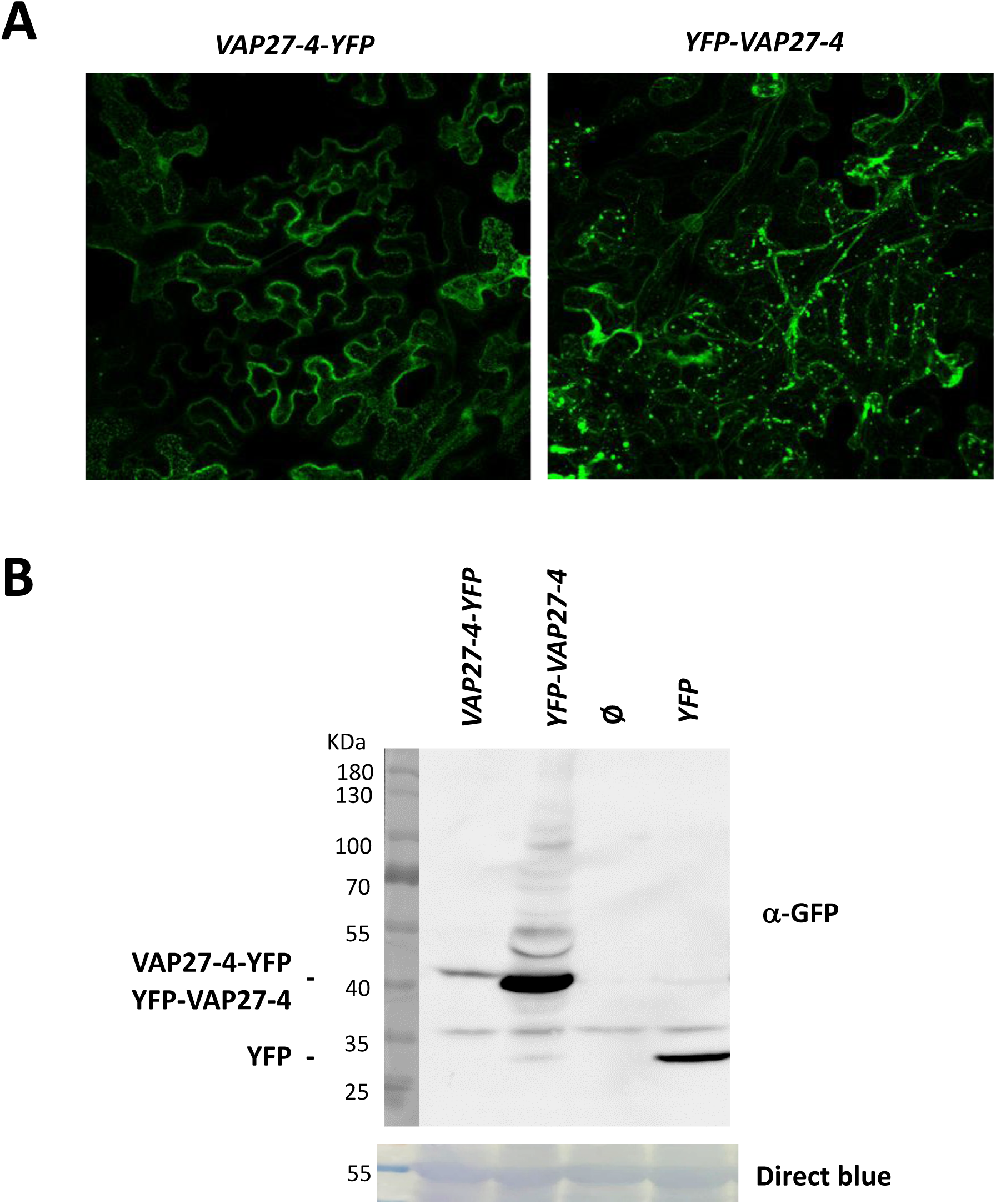
VAP27-4 subcellular localization in *N. benthamiana*. **A.** Representative Z-stack images of *N. benthamiana* leaves agroinfiltrated with *VAP27-4-YFP* or *YFP-VAP27-4* constructs, obtained using fluorescence confocal microscopy. Representative images of three independent experiments are shown. The YFP fluorescence in green is shown. **B.** Western Blot analysis from a total extract of *N. benthamiana* leaves agroinfiltrated with the indicated constructs. The anti-GFP antibody was used for protein detection (upper panel). Direct blue staining of the membrane before antibody incubation is shown at the bottom part as a loading control. Representative images of three independent experiments with three biological replicates are shown. The resulting bands are indicated on the left (theoretical size: VAP27-4-YFP/YFP-VAP27-4 49 kDa; YFP 24.4 kDa).

**Video S1.**
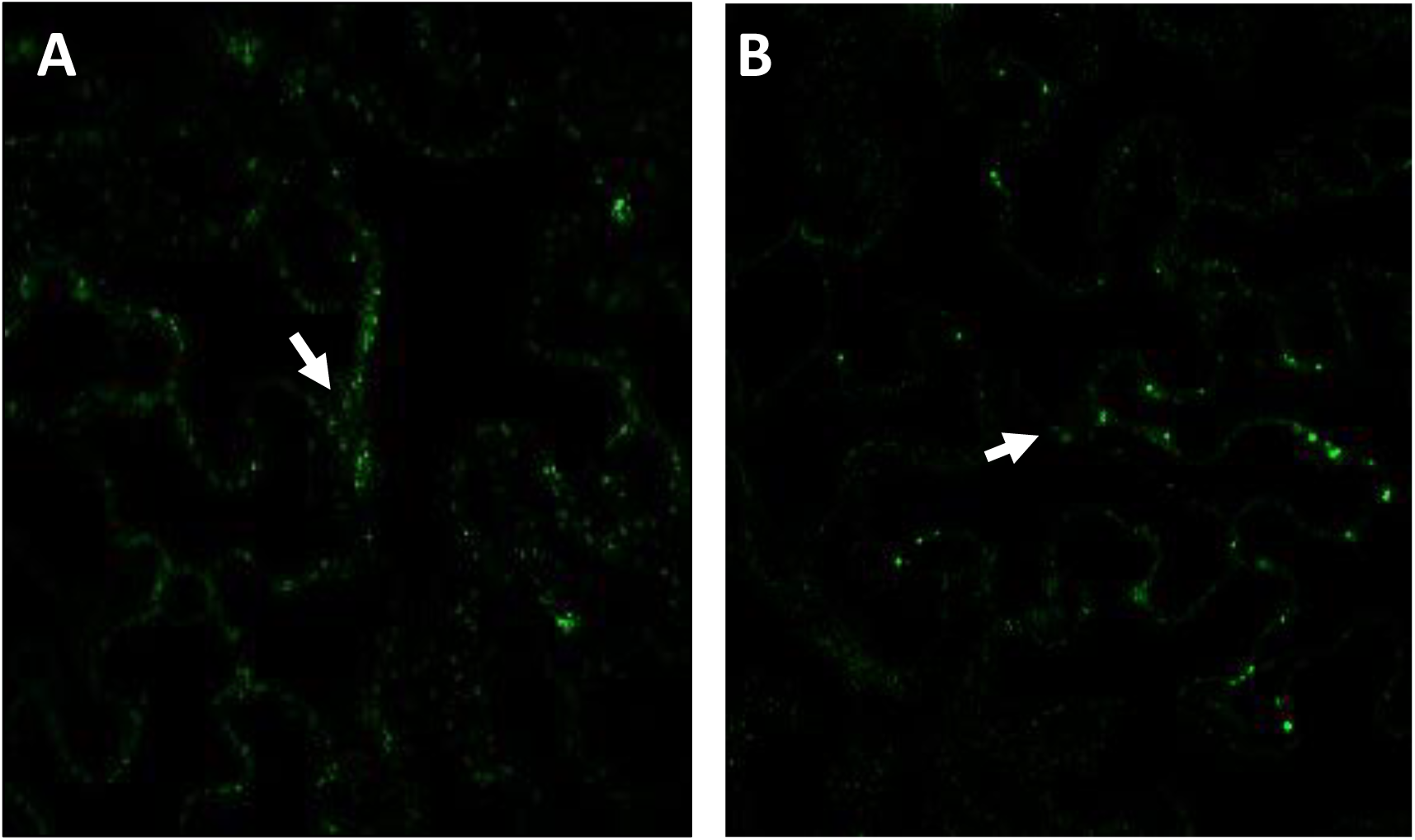
Time-lapse of the VAP27-4 subcellular localization in *N. benthamiana*. **I**mages of representative cells of *N. benthamiana* leaves agroinfiltrated with *YFP-VAP27-4* construct, obtained using fluorescence confocal microscopy (A, 10 frames - 4.37 s; B, 16 frames – 7.29 s). Representative images of three independent experiments are shown. The YFP fluorescence in green is shown. Structures moving from PM to ER **A.** or from ER to PM **B.** are shown. Arrows point to the nucleus in the cell.

**Figure S7.**
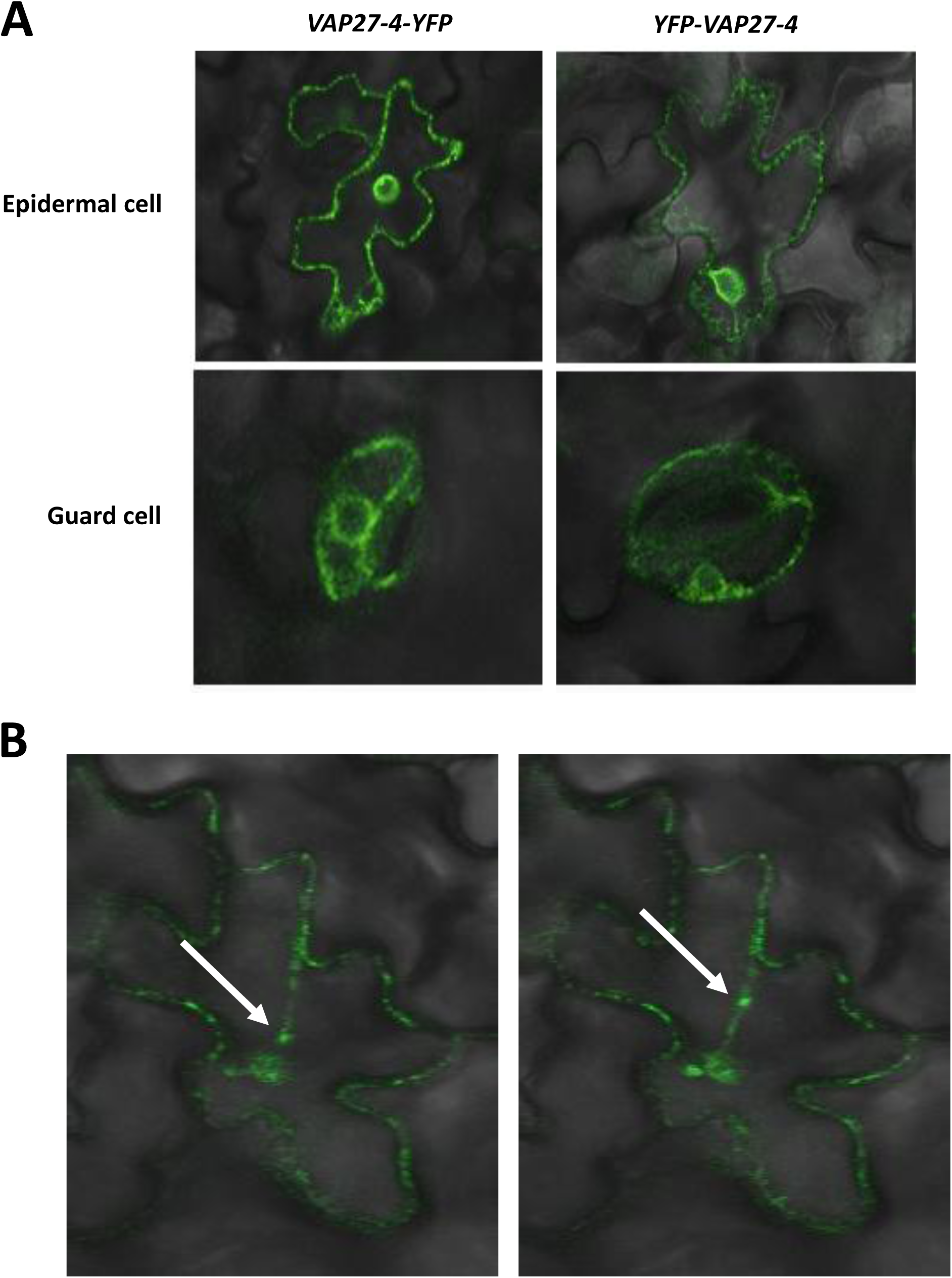
VAP27-4 subcellular localization in Arabidopsis. **A.** Images of representative epidermal and guard cells of leaves of *A. thaliana* lines stably transformed with the *35S::VAP27-4-YFP or 35S::YFP-VAP27-4-YFP* construct, obtained using fluorescence confocal microscopy. Representative images of three independent seedlings are shown. The merge of the YFP fluorescence in green and the bright field is shown. **B.** Images of a representative epidermal cell of leaves of the *A. thaliana* line stably transformed with the *35S::YFP-VAP27-4* construct at different times (0.45 s seconds difference), obtained using fluorescence confocal microscopy. Representative images of three independent seedlings are shown. The YFP fluorescence in green is shown. Arrow, mobile structure.

**Figure S8.**
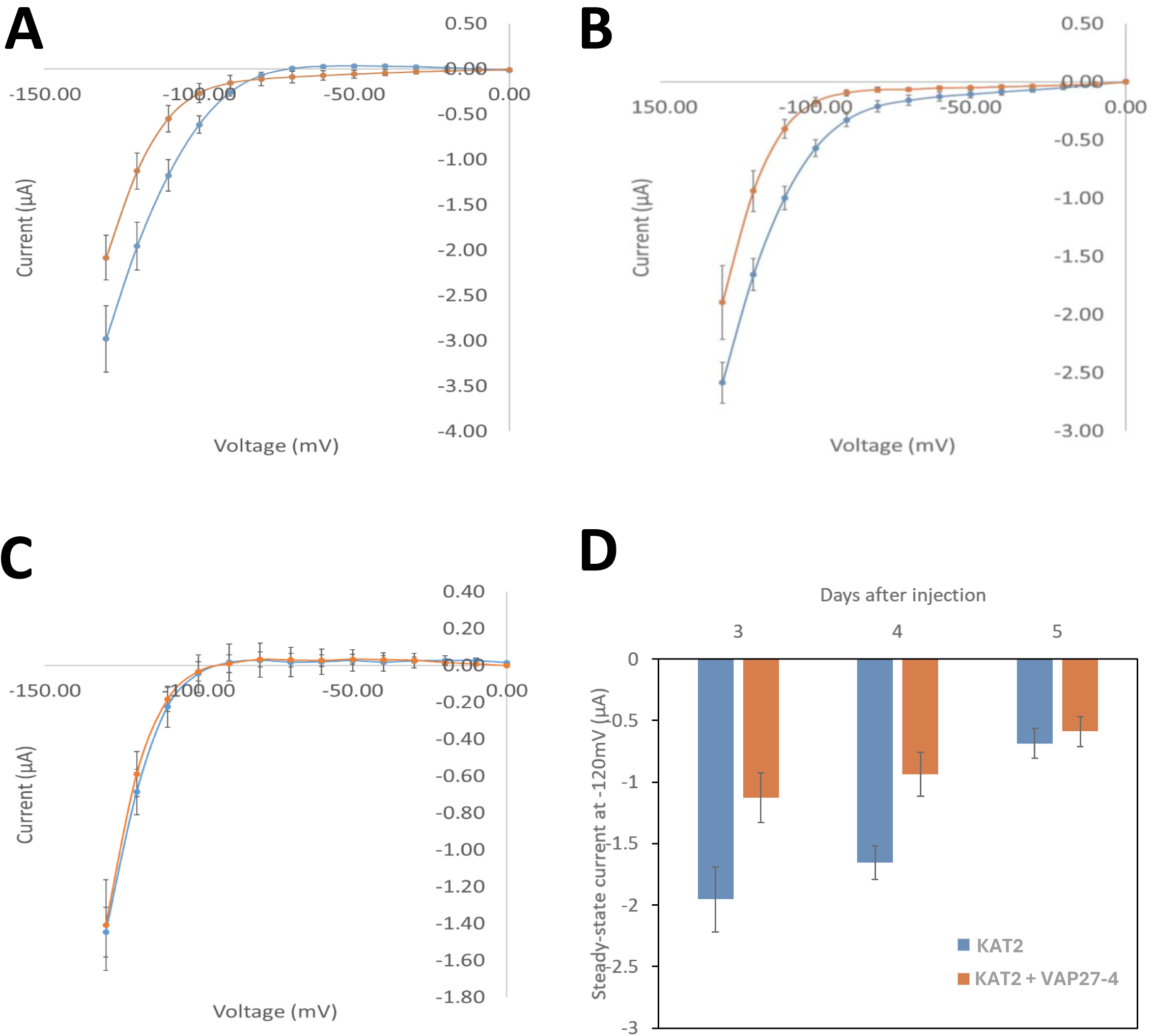
VAP27-4 co-expression decreases KAT2-mediated currents in Xenopus oocytes. **A-C.** KAT2 current (I) –voltage (V) relationships of oocytes co-expressing KAT2 and VAP27-4 (orange line) and expressing KAT2 alone (blue line) at days 3, 4 and 5 after injection. **D**. Average steady-state current values at -120 mV for KAT2 and KAT2+VAP27-4 at 3, 4 and 5 days after injection.

**Figure S9.**
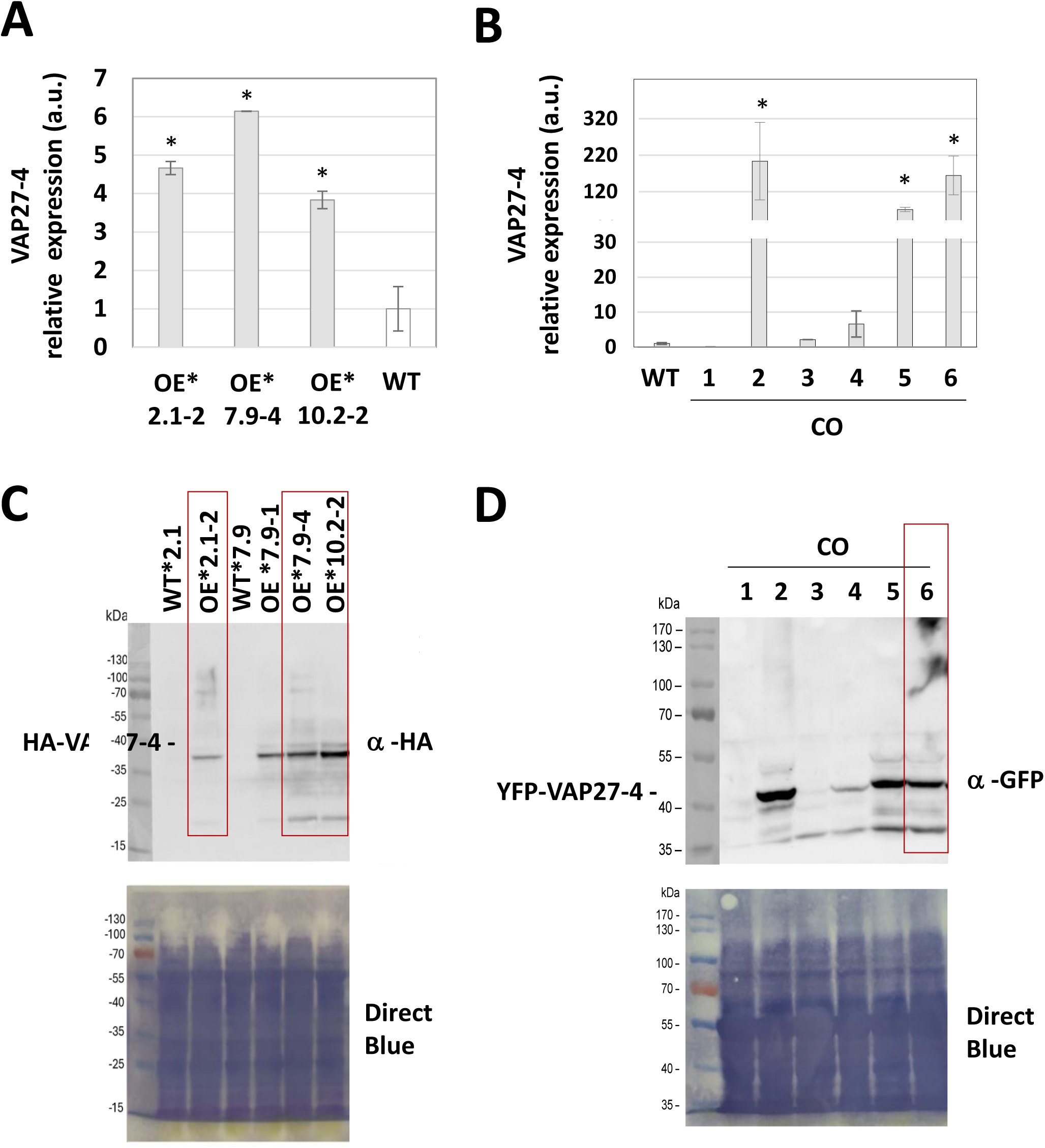
Expression of *VAP27-4* in the overexpressing lines. **A-B.** Total RNA from leaves of adult *A. thaliana* plants grown under long-day conditions was analyzed by RT-qPCR, from WT and two independent *35S::HA-VAP27-4* OE lines, all of them expressing *35S::KAT1-YFP* (*) (A) and six independ*ent vap27-4* + *35S::HA-VAP27-4* (CO) lines (B). Expression values of the *VAP27-4* gene were normalized with respect to the *GADPH* control gene. The WT line was taken as reference (value = 1). The mean of at least two biological replicates is shown with their respective standard deviation. Statistical analysis of each sample with respect to WT: *, p-value ≤ 0.05. a.u., arbitrary units. **C-D.** Western Blot analysis from a total protein extract from leaves of three independent *35S::HA-VAP27-4* OE lines (C) and six independ*ent vap27-4* + *35S::HA-VAP27-4* CO lines (D). The anti-HA (C) and anti-GFP (D) antibodies were used as indicated for protein detection. Direct blue staining of the membrane before antibody incubation is shown in the bottom part as a loading control.

**Figure S10.**
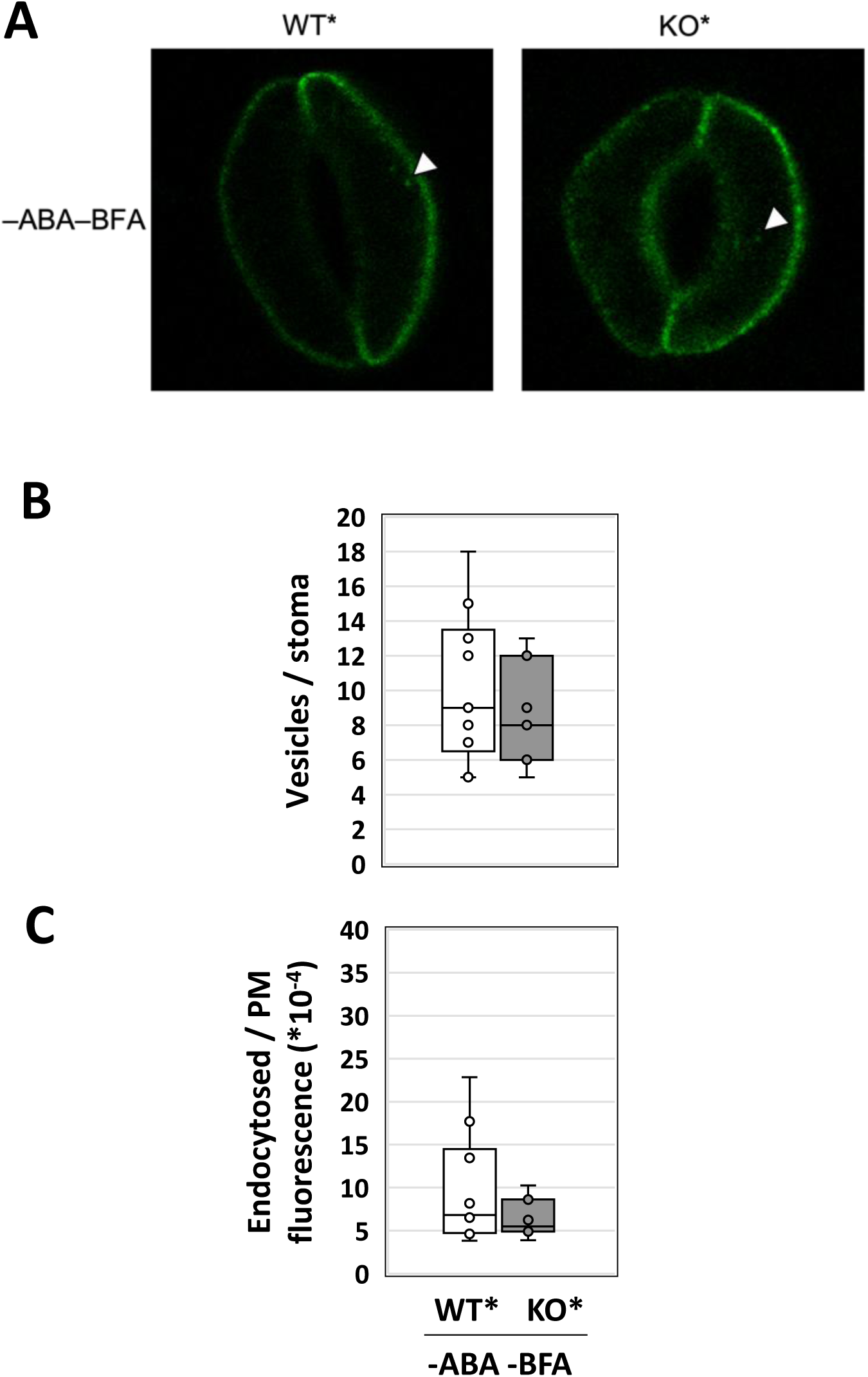
Effect of VAP27-4 on KAT1 endocytosis in Arabidopsis guard cells without ABA. **A.** Representative confocal images of a section of a stoma from leaves of *A. thaliana* WT and *vap27-4* KO line, both of them expressing *35S::KAT1-YFP* (*). Seedlings were grown for 10 long days under control conditions without ABA treatment. The YFP fluorescence in green is shown. White arrowheads point to KAT1-YFP fluorescence within the cell. Representative images of three independent experiments are shown. **B.** Box-whiskers plot of the number of vesicles with KAT1-YFP per stomata, in the whole stomata analyzed in panel A. **C.** Box-whiskers plot of endocytosed KAT1-YFP fluorescence versus KAT1-YFP fluorescence in PM, in the whole stomata analyzed in panel A. Data were obtained from at least three different seedlings, measuring a total of seven stomata or more per line and condition. Statistical analysis of each sample with respect to WT showed not significant differences.

**Figure S11.**
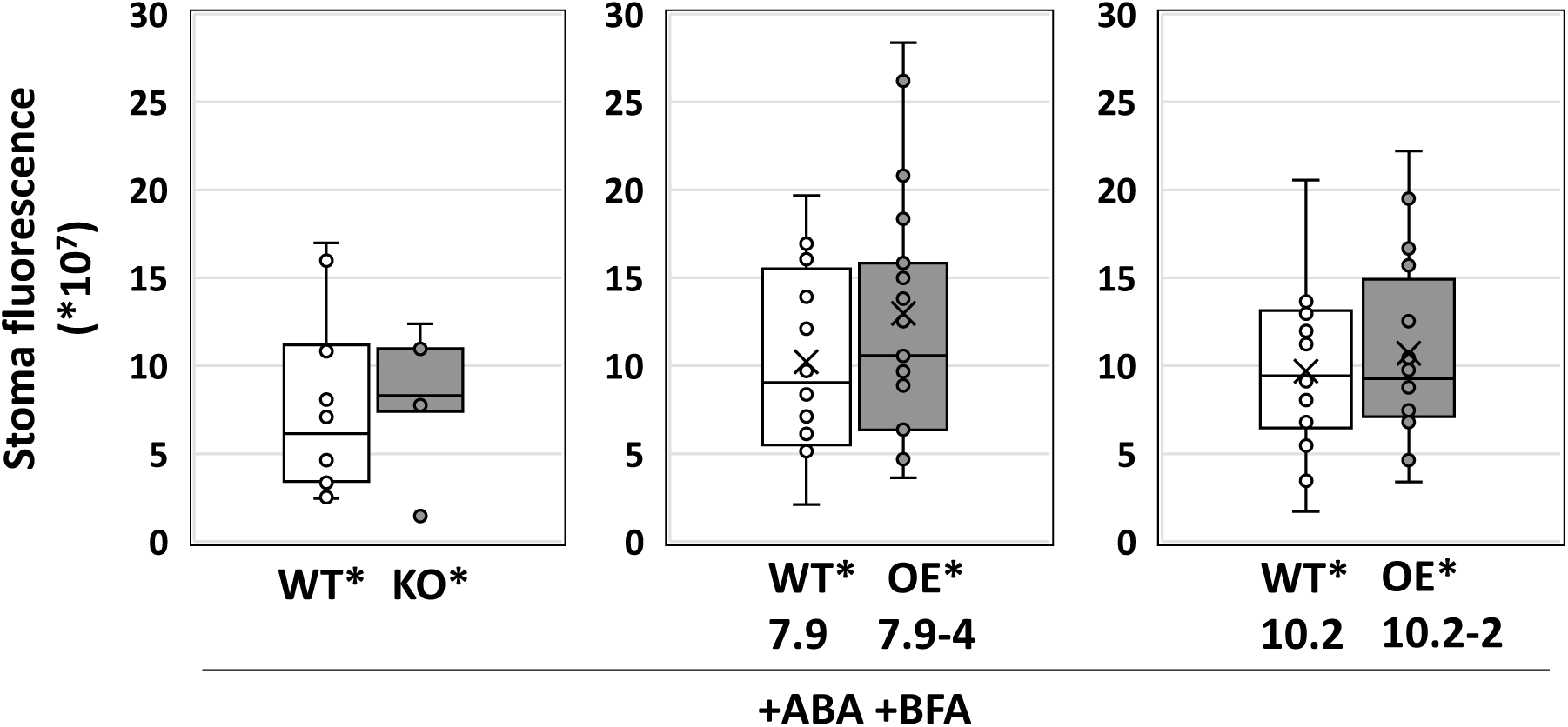
Control for the effect of VAP27-4 on KAT1 endocytosis in Arabidopsis guard cells. Box-whisker plots of the stoma fluorescence, in the whole stomata shown in Figure 6. Data were obtained from at least three different seedlings, measuring a total of seven stomata or more per line and condition. Statistical analysis of each sample with respect to WT showed no significant differences.

**Figure S12.**
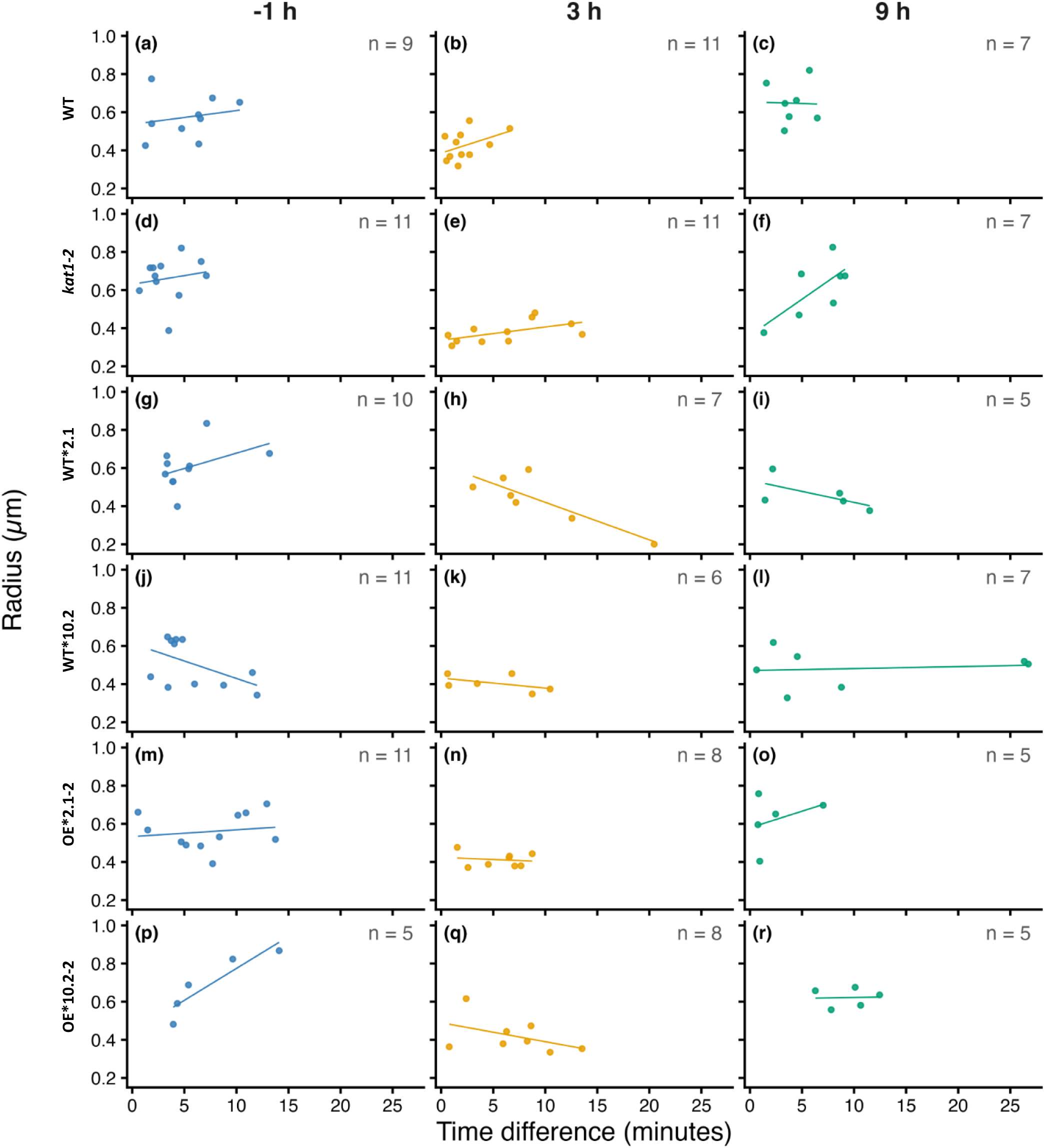
Microbubble radius as a function of time difference between sample exposure to the transmitted light of the microscope optical system and microbubble video capture across genotypes and timepoints. Points represent individual measurements and lines show linear regressions of radius against time difference within each genotype-timepoint combination in Figure 7. None of the 18 fitted slopes differed significantly from zero after Holm adjustment for multiple comparisons (all adjusted p > 0.05) indicating no significant effect of transmitted light exposure on microbubble radius.

## Notes

### Competing Interest Statement

The authors have declared no competing interest.

